# Selective Activation of TASK-3-containing K^+^ Channels Reveals Their Therapeutic Potentials in Analgesia

**DOI:** 10.1101/657387

**Authors:** Ping Liao, Yunguang Qiu, Yiqing Mo, Jie Fu, Zhenpeng Song, Lu Huang, Suwen Bai, Yang Wang, Jia-Jie Zhu, Fuyun Tian, Zhuo Chen, Nanfang Pan, Er-Yi Sun, Linghui Yang, Xi Lan, Yinbin Chen, Dongping Huang, Peihua Sun, Lifen Zhao, Dehua Yang, Weiqiang Lu, Tingting Yang, Junjie Xiao, Wei-Guang Li, Zhaobing Gao, Bing Shen, Qiansen Zhang, Jin Liu, Hualiang Jiang, Ruotian Jiang, Huaiyu Yang

## Abstract

The paucity of selective agonists for TASK-3, a member of two-pore domain K^+^ (K2P) channels, has contributed to our limited understanding of its biological functions. By targeting a novel druggable transmembrane cavity using a structure-based drug design approach, we discovered a biguanide compound, CHET3, as a highly selective allosteric activator for TASK-3-containing K2P channels, including TASK-3 homomer and TASK-3/TASK-1 heteromer. CHET3 displayed unexpectedly potent analgesic effects *in vivo* in a variety of acute and chronic pain models in rodents that could be abolished by pharmacology or genetic ablation of TASK-3. We further found that TASK-3-containing channels anatomically define a unique subset population of small-sized, TRPM8, TRPV1 or tyrosine hydroxylase-positive nociceptive sensory neurons and functionally regulate their membrane excitability, supporting CHET3 analgesia in thermal hyperalgesia and mechanical allodynia under chronic pain. Overall, our proof-of-concept study reveals TASK-3-containing K2P channels as a novel druggable target for treating pain.

**One Sentence Summary:** Identification of a novel drug target and its new hit compounds for developing new-generation non-opioid analgesics.

## INTRODUCTION

Currently available analgesics do not completely treat pain, and some of these analgesics, particularly opioids, provoke social problems (*1*). Thus, discovering new therapeutic targets for developing new-generation analgesics is an urgent need. In particular, targets that treat a variety of pain with similar potency but fewer side effects than μ-opioid receptors are keenly awaited. In this regard, two-pore domain K^+^ (K2P) channels hold great promise (*2*) because they produce background leak K^+^ currents (*3*) and the activation of which in nociceptors theoretically inhibits pain signaling (*4–6*). The expression of TASK-3 (*Kcnk9*), a K2P channel, has been detected in the peripheral and central nervous system (*7, 8*), including in human dorsal root ganglia (*9*). Recent evidence has suggested that TASK-3 is involved in the perception of cold (*10*), and variations in the *Kcnk9* gene are associated with breast pain in breast cancer patients (*11*). However, its functional and anatomical involvement in chronic pain remains largely unknown. More importantly, the paucity of selective agonists limits the drug target validation of TASK-3, leaving the notion that selective activation of TASK-3 alleviates pain remains to be tested experimentally. Here, we sought to discover selective activators for TASK-3 and to use the activators as tool compounds to reveal the translational potentials and the underlying mechanisms of TASK-3 in treating pain.

## RESULTS

### Discovery of the selective activator CHET3 for TASK-3-containing channels

We set out to discover selective activators for TASK-3 via structure-based virtual screening. Since no crystal structure of TASK-3 has yet to be determined, we sought to build a structural model using homology modeling. First, a crystal structure was chosen as the template. To this end, Fpocket 2.0 server (*12*) was applied to detect pockets in the reported crystal structures of K2P channels. In this computation, a druggability score greater than 0.5 (the threshold) means the pocket might be druggable. We found that the cavity under the intracellular side of the filter and the nearby crevice between TM2 and TM4 in four crystal structures (with PDB codes 4RUE, 3UKM, 4XDK and 4XDL) (*13–15*) had druggability scores greater than 0.5 (fig. S1). Thus, this cavity may be a drug binding pocket. Among the four crystal structures, the structure of TREK-2 channel (PDB code 4XDL) is a suitable template for building the structural model of TASK-3 because this structure has good sequence identity (31%) and expectation value (3E-32) in the sequence alignment generated using the BLAST program (blastp algorithm) and the Clustal Omega server (*16, 17*). Moreover, this TREK-2 structure stood out from the template searching results (with the best QSQE value: 0.66) in the SWISS-MODEL server (*18, 19*). Thus, the structural model of TASK-3 was built based on this crystal structure with Modeller (*20*). Then, based on this model, we performed virtual screening targeting the pocket (Fig. 1, A and B) with SPECS and ChemBridge databases. A few hits were selected for the whole-cell patch-clamp electrophysiological tests in HEK-293T cells overexpressing recombinant human TASK-3, which led to the discovery of a biguanide compound CHET3, a novel TASK-3 activator (half-maximum effective concentration (EC50) 1.4 ± 0.2 μM, Fig. 1, C to F). CHET3 maximally enhanced TASK-3-mediated K^+^ currents by ∼4-fold, which could be reversed by washout (Fig. 1D) or with pharmacological blockade by PK-THPP (86 ± 3% inhibition at 0.5 μM, n = 6 cells, representative traces shown in Fig. 1E). PK-THPP is a selective inhibitor for TASK-3 against other K2P and other K^+^ channels including potassium voltage-gated channel subfamily A member 5 (Kv1.5), human ether-à-go-go–related gene (hERG) and G protein-activated inward rectifier potassium channel 4 (KATP) (*21*), and we further showed 0.5 μM PK-THPP was insensitive to voltage-gated K^+^ channel subfamily B member 1 (Kv2.1) and large conductance Ca^2+^-activated K^+^ channel (BK) channels (fig. S2). In single-channel recordings by inside-out patches, CHET3 enhanced the channel openings mediated by TASK-3 (Fig. 1, G to J), further supporting that CHET3 directly activated TASK-3. Increases in open probability were observed in both the inward and outward directions in response to 3 μM CHET3 (Fig. 1J).

**Fig. 1.**
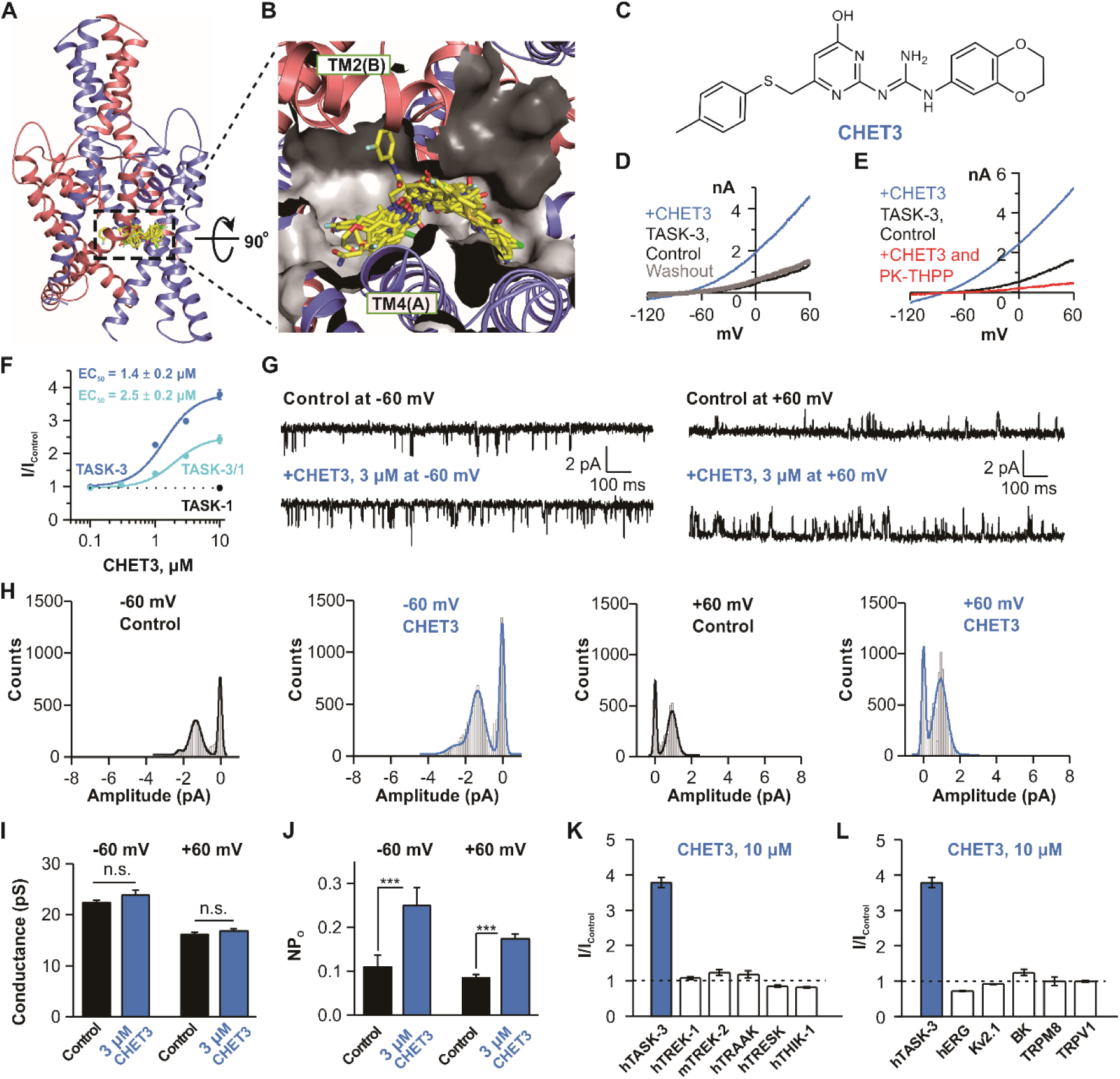
Structure-based ligand discovery of CHET3. (**A**) Pocket in a TASK-3 homology model used for virtual screening. Representative docking poses were shown. **(B)** Cytoplasmic expanded view of the pocket and the docking poses. (**C**) Chemical structure of CHET3. (**D** and **E**) Exemplar whole-cell path-clamp recordings showing the activation of TASK-3 by 10 μM CHET3 and blockade of 0.5 μM PK-THPP. (**F**) CHET3 dose-response curves for TASK-3 (n = 7) and TASK-3/TASK-1 (n = 5). (**G**) Representative single-channel current traces from inside-out patches showing the activation of TASK-3 by CHET3 at −60 mV and +60 mV. (**H**) Histograms of the single-channel currents which were fitted by Gaussian distributions. (**I** and **J**) Analysis of conductance changes and NPo (channel number times open probability) changes from the single-channel currents (n = 9; paired *t* test). (**K**) Summary for the effects of CHET3 on several other K2P channels (n = 7-10). (**L**) Summary for the effects of CHET3 on hERG, Kv2.1, BK, TRPM8 and TRPV1 channels (n = 5-7). Data in (F, I to L) are shown as mean ± SEM. * *P* < 0.05, \*\**P* < 0.01, \*\*\**P* < 0.001.

In addition to forming homomer channels, the TASK-3 subunit can efficiently form heteromer channels with the TASK-1 subunit (*22*). Electrophysiological assays showed that CHET3 could activate TASK-3/TASK-1 (*23, 24*) with an EC50 value of 2.5 ± 0.2 μM with a reduced maximal efficacy of ∼2.4-fold (Fig. 1F), and this activation could also be blocked by PK-THPP (90 ± 2% inhibition at 0.5 μM, n = 5 cells, representative traces shown in fig. S3A). However, CHET3 did not activate TASK-1 channels up to 10 μM (Fig. 1F and fig. S3B). Thus, CHET3 is an activator specific for the TASK-3 homomer and TASK-3/TASK-1 heteromer, two TASK-3-containing channels, with high selectivity against the structurally most related K^+^ channel TASK-1. In the subsequent sections, we use TASK-3-containing channels to represent TASK-3 homomer and TASK-3/TASK-1 heteromer.

Next, we further examined the selectivity of CHET3. Electrophysiological assays of several other K2P channels, including TREK-1, TREK-2, TRAAK, TRESK and THIK-1, further supported that CHET3 has high subtype selectivity among the K2P family (Fig. 1K and fig. S3B). Furthermore, we found that 10 μM CHET3 has high selectivity against hERG, Kv2.1 and BK channels, which are three K^+^ channels sharing similar filter structure and dynamics with K2P (*25*), as well as transient receptor potential cation channel subfamily M member 8 (TRPM8) and transient receptor potential cation channel subfamily V member 1 (TRPV1) (Fig. 1L, fig. S3, C to G).

We also excluded agonizing and antagonizing functions of CHET3 on pain-related G protein-coupled receptors (GPCRs) by testing the cellular function of μ-opioid receptor (μOR), 5-hydroxytryptamine receptor 1B (5-HT_1B_R) and cannabinoid receptor type 1 (CB_1_R) upon treatment with 10 μM CHET3 (fig. S4). Collectively, these results indicate that CHET3 is a selective activator of TASK-3-containing channels.

### Activation mechanism of CHET3

Binding models derived from docking simulation were optimized by molecular dynamics (MD) simulations (fig. S5), which revealed the predominant binding mode of CHET3 within the pocket (Fig. 2, A and B). We next examined the ligand-channel interactions in this binding mode using mutagenesis experiments and RosettaLigand (*26, 27*). Residues T93 and T199 indirectly interacted with CHET3 by water bridges (Fig. 2, A and B). The two residues belong to the filter region, and T93A and T199A mutations led to nonfunctional channels (fig. S6, A and B). F125 may form a π-π interaction with CHET3, and other surrounding residues, including I118, F125, T198, L232, I235, F238 and L239, likely contribute hydrophobic interactions with the ligand. RosettaLigand computations based on this MD mode predicted that the I118A, F125A, L232A, I235A, F238A and L239A mutants decrease CHET3 binding, while the T198A mutant should not (Fig. 2C). Indeed, a saturating concentration of CHET3 (10 μM) showed no activation on F125A, I235A, and F238A mutants and reduced activation on L239A mutant, whereas CHET3 activated T198A similarly to the wild-type (WT) (Fig. 2D). Mutant L232A is nonfunctional (fig. S6C). Although CHET3 did not show reduced activation on I118A (Fig. 2D), the experimental results are generally consistent with the computational predictions. To gain further insight into the action mechanism of CHET3, MD simulations were carried out on *apo* TASK-3 for comparison with CHET3-bound TASK-3 (Fig. 2, E and F). In two out of three independent simulations for the *apo* system, the channel selectivity filter tended to stay in a nonconductive-like conformational state (Fig. 2, E and F). In contrast, in all three simulations for the CHET3-bound system, the channel filter adopted a conductive-like state (Fig. 2, E and F). Furthermore, our simulations supported the previous report by González *et al.* that the H98 residue plays a role in modulating the extracellular ion pathway in TASK-3 (*28*). In the simulations of the CHET3-bound TASK-3, H98 adopted a conformation opening the extracellular ion pathway (Fig. 2E and fig. S7A). In contrast, in ligand-free mode, H98 has a high probability of adopting a conformation closing this pathway (Fig. 2E and fig. S7B).

**Fig. 2.**
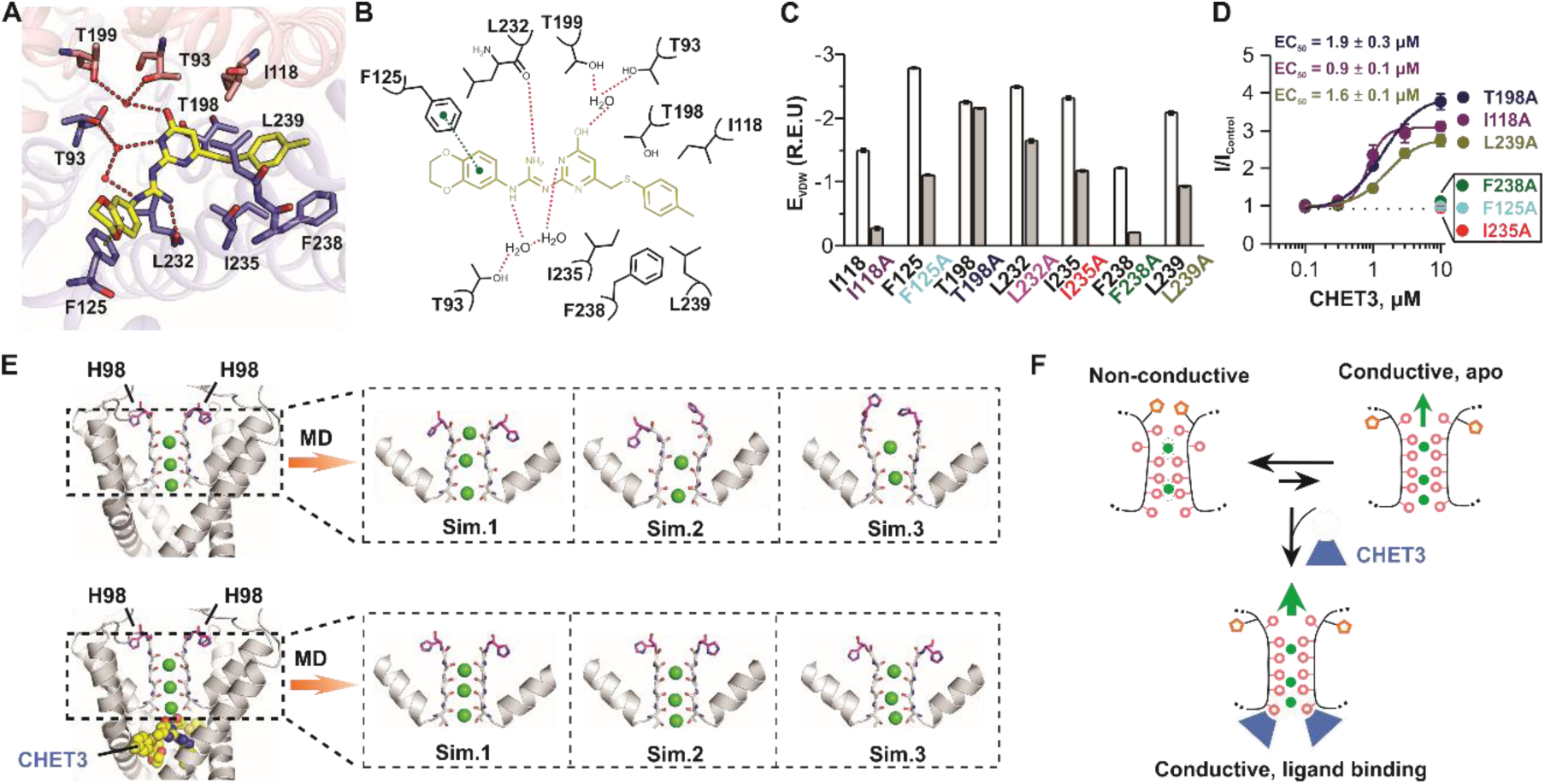
Activation mechanism of CHET3 on TASK-3. (**A** and **B**) 3-dimentional and 2-dimentional diagrams showing interactions between CHET3 and TASK-3. Hydrogen bond (red dash) and π-π interaction (green dash) were shown. (**C**) Computations showing the contributions of seven residues and their mutations to CHET3 binding. Energy unit is Rosetta Energy Unit (R.E.U.). (**D**) Dose-response curves of six mutations on CHET3 activity (n = 6). Data are shown as mean ± SEM. (**E**) Selectivity filter conformations of the *apo* TASK-3 and the CHET3-bound TASK-3 revealed by MD simulations, including bound potassium ions (green spheres), carbonyl oxygen (red sphere) rotation of filter residues, and movements of residue H98 (purple sticks). (**F**) Schematic representation of the action mechanism of CHET3 on TASK-3.

### CHET3-induced analgesia in rodents

Next, we systematically evaluated CHET3 in analgesia. The anti-nociceptive effect of CHET3 was first assessed by the tail immersion test at 52 °C. CHET3 displayed dose-dependent analgesia with a fast onset (30 min) after intraperitoneal (i.p.) injection, with a maximal effect at a dose of 10 mg/kg (Fig. 3A). Hereafter, 10 mg/kg i.p. injection was used for most of the following animal studies. Interestingly, CHET3 was only effective in response to a noxious cold stimulus (5 °C) or a noxious heat stimuli (46 °C and 52 °C) but not to physiological stimuli (20 °C and 40 °C) (Fig. 3A). Importantly, the CHET3 analgesia was fully blocked by the co-administration of PK-THPP, and PK-THPP alone also produced a nociceptive effect in the tail immersion test at 46 °C (Fig. 3A). Next, both the early and late phases (*29*) of acute inflammatory pain induced by formalin were attenuated by CHET3 (Fig. 3B), suggesting at least a peripheral effect of CHET3. The paw pressure test revealed that CHET3 reduced mechanical pain in mice, and the effect was fully blocked by PK-THPP (Fig. 3C). Next, we evaluated the analgesic effects of CHET3 on chronic pathological pain. In the spared nerve injury (SNI)-induced neuropathic pain mouse model, CHET3 significantly reduced the frequency of hind paw lifting (Fig. 3D), an indicator of spontaneous/ongoing pain behavior in the SNI model (*30*). In the cold plantar test, CHET3 attenuated the cold hyperalgesia in SNI in the development (SNI 7 d) and maintenance (SNI 14 d and 21 d) stages of chronic pain, which could be reversed by PK-THPP. PK-THPP alone, however, had no effect in the cold plantar test in SNI mice (Fig. 3E). CHET3 was more effective in relieving cold hyperalgesia than pregabalin, a first-line agent for the treatment of neuropathic pain (Fig. 3F). In SNI mice, CHET3 had little effect on alleviating mechanical allodynia in the von Frey test. However, in SNI rats, 10 mg/kg CHET3 attenuated mechanical allodynia throughout the different stages of chronic pain in the von Frey test, which could be reversed by PK-THPP (Fig. 3G). PK-THPP alone had no effect on pain in the von Frey test in SNI rats (Fig. 3G). The analgesic effect of CHET3 at a dose of 20 mg/kg exhibited a faster onset (30 min post injection) with similar efficacy to 10 mg/kg CHET3 (fig. S8) in the von Frey test in SNI rats. In chronic inflammatory pain induced by the Complete Freund’s Adjuvant (CFA), CHET3 reduced heat hyperalgesia in the Hargreaves test, which was blocked by PK-THPP (Fig. 3H). In addition, PK-THPP injection alone aggravated heat hyperalgesia (Fig. 3H).

**Fig. 3.**
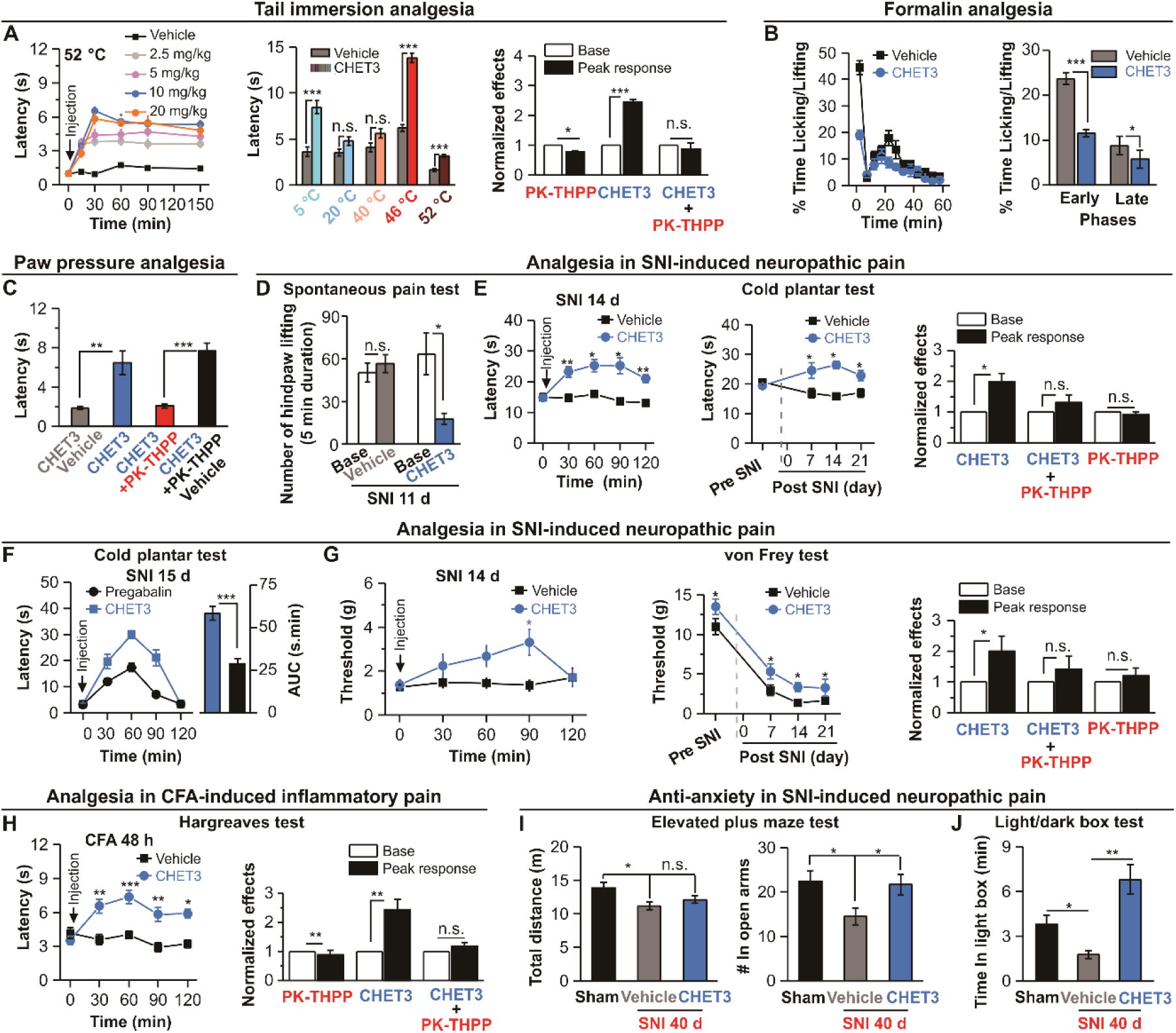
Analgesic effects of CHET3 in rodents. (**A**) *left*, Time profile for dose-dependent analgesia by CHET3 in tail immersion test at 52 °C (n = 6-10); *middle*, CHET3 analgesia in tail immersion tests at different temperatures (n = 9-11; unpaired *t* test); *right*, Summary for PK-THPP effects (n = 8-10; unpaired *t* test). (**B**) Summary for CHET3 analgesia in formalin (2.5%, 20 μl) test (n = 8; unpaired *t* test). (**C**) CHET3 and PK-THPP on the paw withdrawal latency in paw pressure test (measured at 45 min post injection, n = 8-10; paired *t* test). (**D**) CHET3 on spontaneous pain within a 10 min duration in mice (measured at 35 min post injection, n = 6-10; unpaired *t* test). (**E**) *left*, Time profile for CHET3 on cold hyperalgesia in mice (n = 10-11; paired *t* test); *middle*, Summary for CHET3 on cold hyperalgesia at different stages in SNI (n = 9-11; unpaired *t* test); *right*, Summary for PK-THPP (n = 10; paired *t* test). (**F**) Comparison of CHET3 and Pregabalin (30 mg/kg, i.p.) in cold plantar test in mice (n =11-12; unpaired *t* test). (**G**) *left*, Time profile for CHET3 in mechanical allodynia in rats (n = 8-13; paired *t* test); *middle*, Summary for CHET3 on mechanical allodynia at different stages in SNI rats (n = 8-13; paired *t* test); *right*, Summary for PK-THPP effects (n = 8; paired *t* test). (**H**) *left*, Time profile for CHET3 on heat hyperalgesia (n = 10-11; paired *t* test); *right*, Summary for PK-THPP effects (n = 11; paired *t* test). (**I** and **J**) CHET3 on anxiety-like behaviors in elevated plus maze test (I) (n = 8-9; unpaired *t* test) and in light/dark box tested (J) (n = 6-8; unpaired *t* test). CHET3 (10 mg/kg) and PK-THPP (15 mg/kg) were administrated via i.p. injections unless specified. Data are shown as mean ± SEM. * *P* < 0.05, \*\**P* < 0.01, \*\*\**P* < 0.001. n.s., not significant.

Chronic pain may induce anxiety (*31, 32*). Compared with Sham mice, SNI mice spent less time in open arms in the elevated plus maze test and spent less time in the light box in the light/dark box test, suggesting anxiety-like behaviors in the SNI mice. The administration of CHET3 30 min before the test significantly alleviated anxiety-like behaviors in both tests (Fig. 3, I and J). Together, our data suggest that CHET3 potently and efficaciously attenuated acute and chronic pain and pain-associated anxiety in rodents, and the analgesic effects of CHET3 could be pharmacologically blocked by the TASK-3 blocker PK-THPP. Importantly, CHET3 had no effects in grip strength, rotarod and open field tests (fig. S9, A to C), suggesting that CHET3 had no effect on the locomotion activities in mice. Since TASK-3 was found to be expressed in mouse carotid boy type-1 cells (*33*), we also evaluated the possible side effects of CHET3 on cardiovascular function in mice or rats. We monitored blood pressure and heart functions using echocardiography, and we did not observe any significant change in blood pressure (fig. S9, D to F) or heart functions including Ejection Fraction (EF) and Fractional Shortening (FS) (table S1) in a post-injection time window between 45 min–90 min, during which CHET3-induced analgesia peaked in most cases. We also monitored the change in body temperature following CHET3 systemic administration, and no significant hyperthermia or hypothermia was observed (fig. S9G).

### Further on-target validation using chemical and genetic approaches

Was CHET3 truly targeting TASK-3 containing K^+^ channels as an analgesic? We next performed additional target validation experiments using chemical and genetic approaches. Medicinal chemistry yielded CHET3-1 and CHET3-2 (Fig. 4A), two derivatives of CHET3. In the CHET3-TASK-3 binding model (Fig. 2, A and B), the dioxane ring may form a π–π interaction with the F125 residue. CHET3-1, in which the dioxane ring is replaced with an aromatic ring, should maintain the π–π interaction. CHET3-2 should lose the π–π interaction since the dioxane ring is replaced by a steric bulk *tert*-butyl group. Binding energy computations based on the binding model suggested that the binding affinity of CHET3-1 to TASK-3 was similar to that of CHET3, while that of CHET3-2 decreased (fig. S10). In accordance, CHET3-1 activated TASK-3 with an EC50 value of 0.5 ± 0.1 μM, while CHET3-2 was inactive (Fig. 4B), further supporting the putative binding model. We reasoned that CHET3-1 should be bioactive in analgesia, whereas CHET3-2 should not, if CHET3 truly targets TASK-3-containing channels to act as an analgesic. Indeed, CHET3-1 attenuated cold hyperalgesia in SNI mice (Fig. 4C), mechanical allodynia in SNI rats (Fig. 4D), and heat hyperalgesia in CFA mice (Fig. 4E), and all of these effects could be reversed by PK-THPP (fig. S11). In contrast, CHET3-2 was completely inactive in all the experiments above (Fig. 4, C to E).

**Fig. 4.**
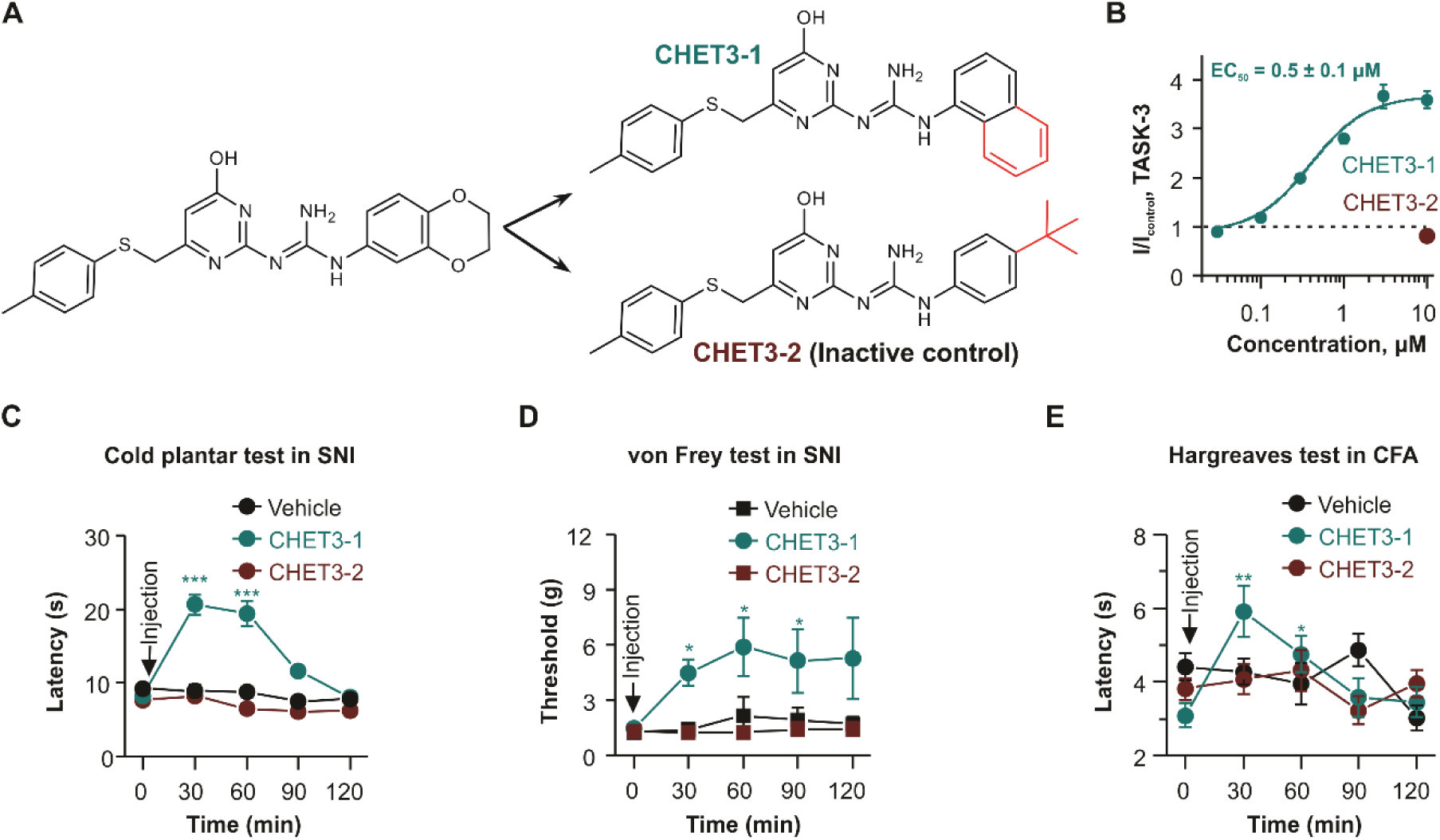
Validation of TASK-3 as analgesic target by using CHET3 derivatives. (**A**) Chemical structures of CHET3 derivatives. (**B**) Dose-response relationships for CHET3-1 (n = 6) and CHET3-2 (n = 7) on TASK-3. (**C** to **E**) CHET3-1 and CHET3-2 in cold hyperalgesia (C) (n = 9-11; paired *t* test), mechanical allodynia (D) (n = 7-8; paired *t* test), and heat hyperalgesia (E) (n = 9-10; paired *t* test). Data are shown as mean ± SEM. \**P* < 0.05, \*\**P* < 0.01, \*\*\**P* < 0.001.

We also generated *Kcnk9* gene knockout (TASK-3 KO) mice (fig. S12). Knocking out *Kcnk9* should abolish the function of TASK-3 homomer and TASK-3/TASK-1 heteromer *in vivo*. In TASK-3 KO mice and WT control mice, we measured the basal sensitivity to nociception, thermal hyperalgesia and mechanical allodynia, and we also evaluated the analgesic effect of CHET3 in these mice. Tail immersion (Fig. 5, A and B), paw pressure tests (Fig. 5C) and von Frey tests in naïve animals (Pre SNI, Fig. 5D) did not reveal any significant difference in baseline nociceptive sensitivity between TASK-3 KO and WT; however, cold plantar (Pre SNI, Fig. 5F) and Hargreaves tests (Pre CFA, Fig. 5G) revealed increased nociceptive cold and heat sensitivity in TASK-3 KO mice. Furthermore, in the chronic pain models, von Frey, cold plantar and Hargreaves tests revealed that TASK-3 KO mice exhibited aggravated mechanical allodynia (Fig 5D), spontaneous neuropathic pain behavior (Fig. 5E), and thermal hyperalgesia (Fig. 5, F and G). As expected, CHET3 was completely inactive in TASK-3 KO mice in all the tests described above (Fig. 5, A to G). Thus, using tool compounds (Fig. 4) and mouse genetics (Fig. 5), we provide strong evidence showing that CHET3 targets TASK-3-containing channels, thus acting as an analgesic, and the loss of TASK-3 contributed to the generation/maintenance of both acute and chronic pain.

**Fig. 5.**
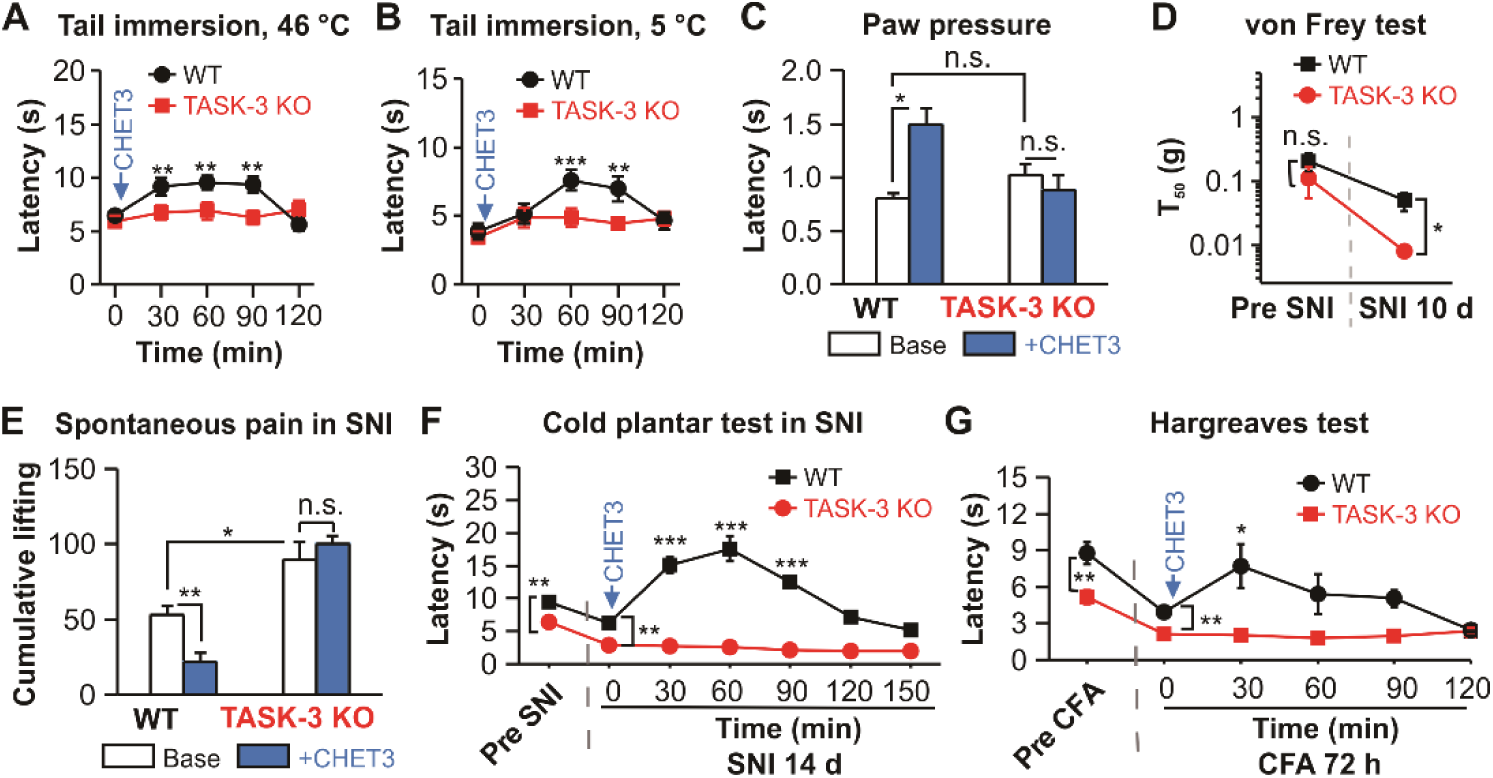
Effects of systemic administration of CHET3 on TASK-3 KO mice. (**A** and **B**) CHET3 had no analgesic effects in tail immersion tests in TASK-3 KO mice (n = 9-12; paired *t* test). (**C**) CHET3 had no analgesic effect in paw pressure tests in TASK-3 KO mice (n = 8 for KO, n = 7 for WT; paired *t* test). Note that no change of baseline sensitivity in nociception for TASK-3 KO mice in A to C. (**D**) TASK-3 KO mice in SNI model exhibited enhanced mechanical allodynia (up-down method, n = 8 for KO, n = 10 for WT; unpaired *t* test). (**E**) TASK-3 KO mice in SNI model exhibited enhanced spontaneous pain activities which was insensitive to CHET3 (n = 6-7; paired *t* test). (**F** and **G**) CHET3 had no analgesic effect in cold plantar test (n = 6-7; paired *t* test) in SNI model and Hargreaves test (n = 8-12; paired *t* test) in CFA model in TASK-3 KO mice. Note that TASK-3 KO mice had shorter paw withdraw latencies in both tests (unpaired *t* test) in base conditions. Data are shown as mean ± SEM. * *P* < 0.05, \*\**P* < 0.01, \*\*\**P* < 0.001. n.s., not significant.

### Distribution of TASK-3-containing channels in sensory neurons

Our pharmacokinetic profile of CHET3 (tables S2 and S3) showed a negligible brain concentration of CHET3 and a high concentration of CHET3 in the plasma in both the naïve and SNI 7-d mice, suggesting that CHET3 mainly acted peripherally. The peripheral effect of CHET3 is also supported by the fact that CHET3 attenuated the early phase of formalin-induced pain (Fig. 3B). These findings, along with the previous finding that TASK-3 in dorsal root ganglion (DRG) neurons mediates cold perception (*10*), strongly suggest that peripheral TASK-3-containing channels contribute largely, if not entirely, to the analgesic effects of CHET3. Therefore, we evaluated the TASK-3 functions/distributions in the peripheral nervous system, particularly in DRG.

We used fluorescence in situ hybridization (ISH) (RNAscope technique) to map the mRNA expression of TASK-3 in DRG and trigeminal ganglia (TG). The specificity of the fluorescent signals was validated by a positive control probe and a negative control probe (see the Methods). *Kcnk9* was identified in a subset of neurons (∼7% of total neurons) in DRG, predominantly in small-sized neurons (diameter ≤ 20 μm) (Fig. 6, A and C), indicative of its specific expression in nociceptors. Interestingly, a much higher expression level of *Kcnk9* (∼14% of total neurons) was found in TG (Fig. 6, A and B). In DRG, approximately 95% of *Kcnk9*^+^ neurons express the TASK-1 subunit, suggesting possible formation of the TASK-3/TASK-1 heteromer in DRG, and approximately 50% of *Kcnk9*^+^ neurons express TRPV1, a well-known noxious heat sensor predominantly expressed in peptidergic nociceptive sensory neurons (*34*). More than 95% of *Kcnk9*^+^ neurons express TRPM8, and very little *Kcnk9*^+^ neurons express TRPA1, two well-known noxious cold sensors (*34*). Furthermore, approximately 50% of *Kcnk9*^+^ neurons express tyrosine hydroxylase (TH), a marker for c-low threshold mechanoreceptors (c-LTMRs) predominantly found in nonpeptidergic nociceptors (*35*), whereas *Kcnk9* rarely colocalizes with *P2rx3* (P2X3), which labels mainly TH^−^ negative, IB4^+^ nonpeptidergic nociceptors (*36*), nor does *Kcnk9* colocalize with *Ntrk2* (TrkB), a marker for Aδ-LTMRs (*35*). Thus, TASK-3 marks a unique subpopulation of both peptidergic and nonpeptidergic nociceptive sensory neurons enriched in thermal sensors (TRPV1, TRPM8) or mechanoreceptors (TH^+^ c-LTMRs) (Fig. 6, D to F), in line with its functional involvement in thermal and mechanical sensation *in vivo*. In agreement with a previous study (*37*), we found that *Kcnk9* expression was downregulated in SNI mice and CFA mice (fig. S13), further suggesting that the downregulation of TASK-3-containing channels contributes to the generation/maintenance of chronic pain.

**Fig 6.**
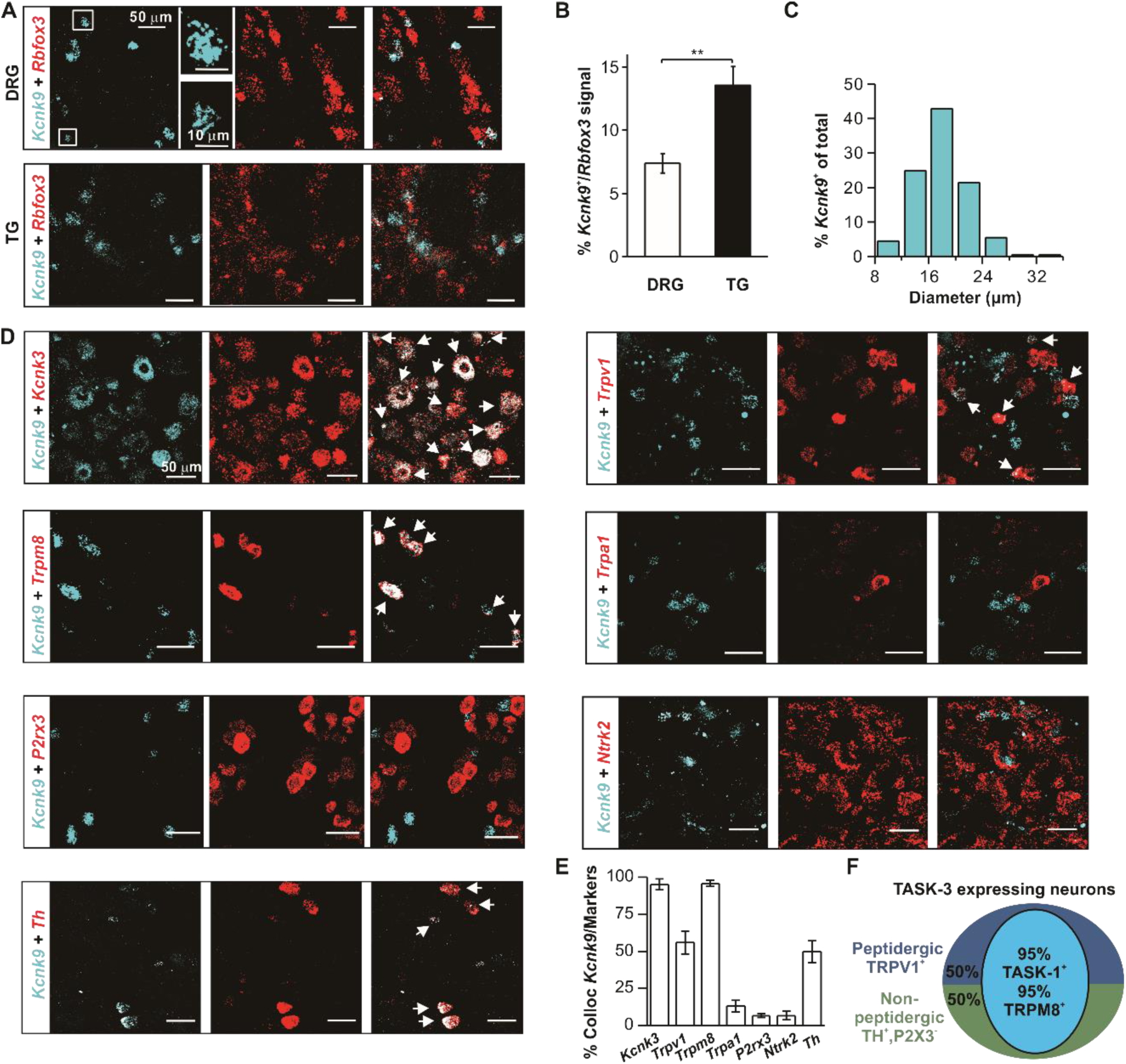
Distribution of TASK-3 in DRG neurons. (**A** and **B**) Images and quantifications showing *Kcnk9* expression in sensory neurons using RNAscope (n = 7 sections from 3 mice) and TG (n = 8 sections from 3 mice) (Mann-Whitney test). Data in (B) are shown as mean ± SEM. (**C**) Quantification of the cell sizes of *Kcnk9^+^* neurons (n = 6 sections from 3 mice). (**D**) Representative images showing *Kcnk9*^+^ neurons expression in different subtype of DRG neurons using RNAscope. (**E**) Bar graph summary for experiments in (D) (n = 4-9 DRG sections from 3-8 mice for each condition). Data are shown as mean ± SEM. (**F**) Schematic summary for the distribution of *Kcnk9*^+^ neurons in DRG. \*\**P* < 0.01.

### Functional roles of TASK-3-containing channels in nociceptive neurons

The functional roles of TASK-3-containing channels were examined by whole-cell patch-clamp recordings in dissociated DRG neurons. Recordings were focused on small DRG neurons (diameter of ∼20 μm, cell capacitance of ∼30 pF) based on the ISH data. To isolate K^+^ currents, voltage ramps from −120 mV to −30 mV were applied. In total, 89 cells were recorded, and 16 cells responded to CHET3 (20.3 ± 6.3%, 11 mice). In the CHET3-sensitive cells, CHET3 enhanced the whole-cell current density by approximately 18%, which could be further inhibited by PK-THPP by approximately 38% at −30 mV (Fig. 7A). We subtracted the CHET3-sensitive current, and we found this current was strongly outwardly rectifying and was tiny between −120 mV and −60 mV, leaving the reversal potential of the CHET3-sensitive current difficult to resolve (Fig. 7B). We further sought to isolate current carried by TASK-3-containing channels by subtracting PK-THPP-sensitive current. We consistently found a similar profile for PK-THPP-sensitive current (fig. S14A), further suggesting the low basal conductance at hyperpolarized membrane potentials, while strong outwardly rectifying represents an intrinsic property of K^+^ currents mediated by TASK-3-containing channels in DRG under our experimental conditions. To increase the drive force of the K^+^ currents in the hyperpolarized potentials, we increased the extracellular K^+^ concentration to 143 mM. Under this condition, the CHET3-sensitive current was reversed at approximately 6.7 mV, which was close to the theoretical value of 1.5 mV for K^+^ conductance (Fig. 7B).

**Fig 7.**
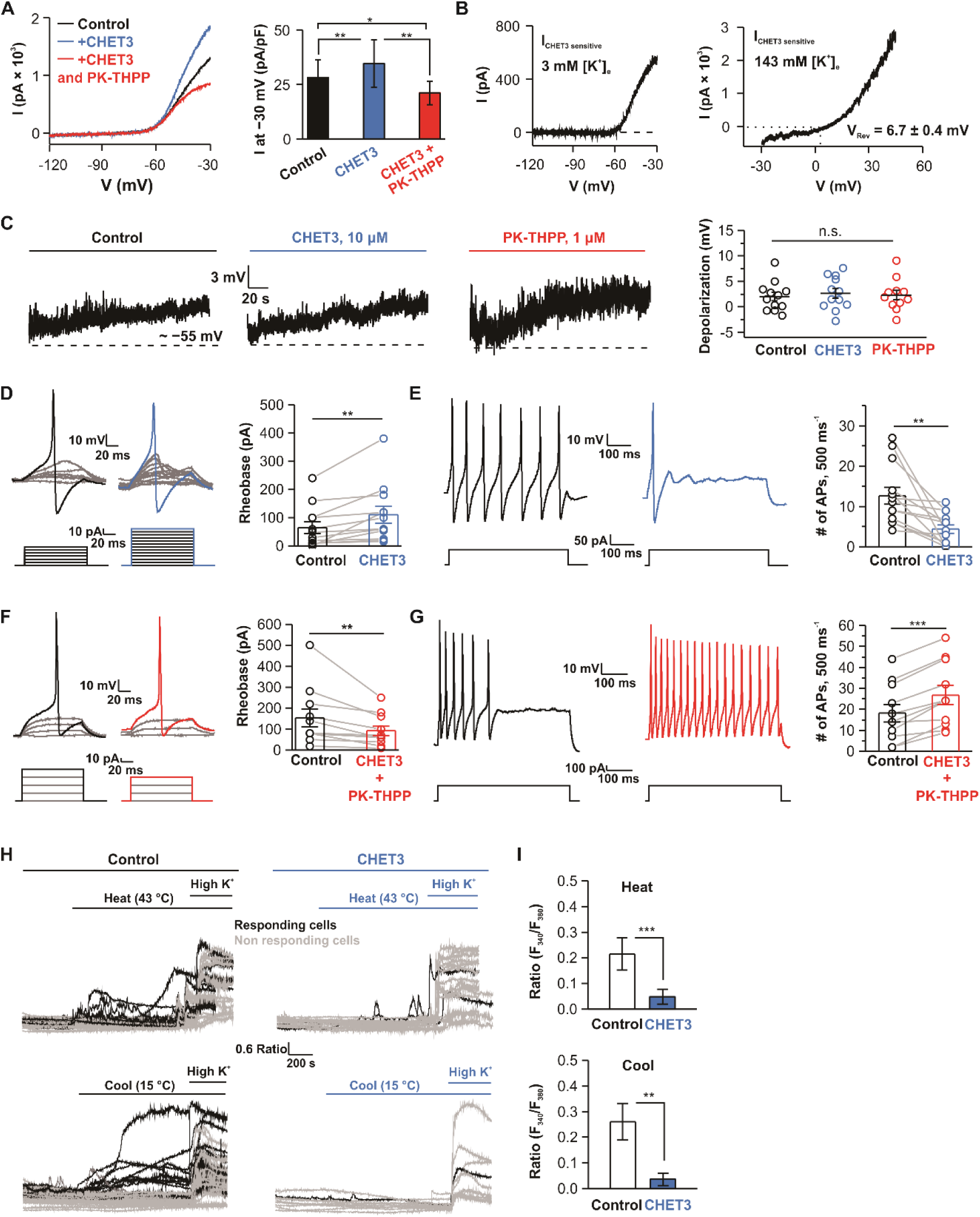
Functional roles of TASK-3 in nociceptive neurons. (**A**) *left*, Representative electrophysiological traces showing CHET3 (10 μM) and PK-THPP (1 μM) effects on K^+^ currents in DRG neurons; *right*, Bar graph summary for experiments in *left* (n = 9 cells in 6 mice; paired *t* test). (**B**) Representative traces showing CHET3-sensitive currents at different extracellular K^+^ concentrations (VRev was determined from n = 5 cells in 3 mice). (**C**) Representative traces and scatter plots showing resting membrane potential (RMP) changes in response to Vehicle, CHET3 or PK-THPP (n = 12 cells for Control and CHET3, n = 11 cells for PK-THPP in 5 mice for each condition; one-way ANOVA test). (**D** and **E**) Traces and bar graph showing CHET3 effect on rheobase and firing frequency in nociceptive neurons (n = 12 cells in 5 mice; paired sample Wilcoxon signed rank test in (D), paired *t* test in (E)). (**F** and **G**) Traces and bar graph showing co-application of CHET3 and PK-THPP on rheobase and firing frequency in nociceptive neurons (n = 11 cells in 3 mice; paired sample Wilcoxon signed rank test in (F), paired *t* test in (G)). (**H**) Individual Ca^2+^ imaging traces from small-sized DRG neurons in representative field of views in response to heat (25 °C-43 °C), cooling (37 °C-15 °C). (**I**) Bar graphs summary for experiments in (H) (Heat: n = 143 cells in 11 coverslips for control, n = 99 cells in 9 coverslips for CHET3; Mann-Whitney test. Cool: n = 87 cells in 9 coverslips for control, n = 46 cells in 9 coverslips for CHET3; Mann-Whitney test. Both experiments were from 3 independent preparations from 6 mice). Data are shown as mean ± SEM. * *P* < 0.05, \*\**P* < 0.01, \*\*\**P* < 0.001. n.s., not significant.

The tiny CHET-3- or PK-THPP-mediated currents at approximately −60 mV suggest that the basal activity of TASK-3-containing channels around the resting membrane potential (RMP) range was low and, thus, that CHET3 or PK-THPP is unlikely to be able to regulate the RMP. To systematically evaluate the regulatory role of CHET3 on the excitability of nociceptive neurons, we first applied a cocktail solution containing menthol and capsaicin, two agonists for TRPM8 and TRPV1 (*38, 39*), respectively, to better identify the nociceptive neurons that likely express TASK-3-containing channels. Only neurons responding to the cocktail (fig. S14B) were studied in the subsequent experiments. Consistent with the low activity of TASK-3-containing channels at approximately −60 mV, the application of CHET3 or PK-THPP or vehicle (Control) did not hyperpolarize the RMP; rather, they all slightly depolarized the membrane by ∼2 mV with no significant difference among the three groups, suggesting that CHET3 or PK-THPP had no specific roles in altering RMP (Fig. 7C). Next, we explored how CHET3 regulates action potentials. In 12 out of 27 neurons, the application of CHET3 markedly increased the rheobase required to elicit the action potentials (APs) by ∼70% and decreased the frequency of APs evoked by suprathreshold current injections by ∼65% (Fig. 7, D and E). In the other 15 cells, CHET3 had no effect on the rheobase and slightly increased the frequency of APs evoked by suprathreshold current injections by 10% (fig. S14, C and D). In 7 of these 12 CHET3-sensitive cells, we were able to further apply PK-THPP, which reversed the effects of CHET3 (fig. S14, E and F). Furthermore, in another independent set of experiments, we coapplied CHET3 and PK-THPP in naïve cells. In 11 out of 27 cells, the coapplication of CHET3 and PK-THPP markedly decreased the rheobase by ∼40% and increased the frequency of APs evoked by suprathreshold current injections by ∼50% (Fig. 7, F and G); in the other 16 cells, coapplication of CHET3 and PK-THPP slightly increased the rheobase by ∼20% but had no effect on the AP frequency evoked by suprathreshold current injections (fig. S14, G and H). Collectively, our electrophysiological data suggest the functional presence of K^+^ currents mediated by TASK-3-containing channels, the enhancement of which reduces the excitability of nociceptive neurons without affecting the RMP.

Finally, Ca^2+^ imaging experiments were performed in acutely dissociated DRG neurons to measure how the activation of TASK-3-containing channels contributes to the thermal sensitivity of DRG neurons. Thermal stimulations elicited Ca^2+^ signals in a portion of small-sized DRG neurons (Fig. 7H, cells with an F_340_/F_380_ ratio ≥ 0.2 were considered responding cells shown in black, and those with an F_340_/F_380_ ratio < 0.2 were considered nonresponding cells shown in gray). We confirmed that these Ca^2+^ signals were temperature-dependent and were mediated by TRP channels because the heat-induced responses could be blocked by 5 μM AMG9810 (TRPV1 antagonist) (*40*) (fig. S15, A and B), and the cool-induced responses could be blocked by 10 μM BCTC (TRPM8 antagonist) (*38*) and 20 μM HC030031 (TRPA1 blocker) (*41*) (fig. S15, A and B). Bath application of 10 μM CHET3 significantly and markedly inhibited the Ca^2+^ signals evoked by cool- or heat-stimulation in small-sized DRG neurons (Fig. 7, H and I), suggesting that the activation of TASK-3-containing channels was able to lower the excitability of the nociceptive neurons in response to external thermal stimulations.

## DISCUSSION

The current study has three major findings: First, we discovered selective agonists for TASK-3-containing channels by targeting a transmembrane cavity under the selectivity filter using structure-based approaches. Second, *in vivo* activation of peripheral TASK-3-containing channels displayed potent analgesia, suggesting a TASK-3-based therapeutic strategy for treating chronic inflammatory and neuropathic pain. Third, our anatomical and functional data highlight the roles of peripheral TASK-3-containing channels in controlling the excitability of nociceptive neurons.

Very recently, Schewe *et al.* reported a class of negatively charged activators (NCAs) that could activate K2P channel, hERG channel and BK channel and revealed that the site below the selectivity filter is the binding site of the NCAs (*25*). In the present work, our virtual screening obtained CHET3, a non-charged compound that acts on this site, further supporting the finding that the site below the selectivity filter is a ligand binding site. It is noteworthy that NCAs are nonselective activators for a variety of K^+^ channels, while CHET3 is highly selective for TASK-3-containing channels, suggesting the versatility of this binding site. Additionally, NCAs and CHET3 may share some common activation mechanisms on K2P channels, as they both influence the conformation of the selectivity filter. Notably, the activation mechanism we describe in this study does not fully explain the selectivity of CHET3. In particular, TASK-1 and TASK-3 are the closest relatives to each other, and the residues below the selectivity filter as well as H98 are conserved. Further studies to elucidate the differential responses of TASK-1 and TASK-3 to CHET3 may be helpful for understanding the selective modulation principle in K2P.

In most cases, the initial proof-of-concept identification of a protein as a potential target is dependent on genetic methods. However, genetic deletion may induce modifications to other genes. This off-target genetic side effect discredits target validation work. This is particularly the case in the field of pain medicine: genetically mutated mice, e.g., Na_v_1.7-null mice and humans exhibited remarkable insensitivity to pain, whereas potent selective antagonists have weak analgesic activity (*42, 43*). Another example related to the K2P field is that migraine-associated TRESK mutations lead to the inhibition of TREK-1 and TREK-2 through frame shift mutation-induced alternative translation initiation (fsATI) to increase sensory neuron excitability and are linked to migraine (*44*). Using chemical probes to validate targets pave another way for later translational research. Regarding in *vivo* applications of chemical probes in target identification and validation, a major issue is whether the observed phenotypes are indeed relevant to the on-target of the probes. In this study, we provide three independent lines of evidence showing that CHET3 targets TASK-3-containing channels to act as an analgesic. First, the TASK-3 inhibitor PK-THPP could block CHET-3-induced analgesia. Second, two structurally similar analogs were discovered and used in the *in vivo* tests. CHET3-1, a TASK-3 activator structurally similar to CHET3, is bioactive in analgesia, and could also be blocked by PK-THPP. CHET3-2, another analog that is highly structurally similar to CHET3, did not activate TASK-3 and was completely ineffective in all the analgesia tests. Finally, CHET3 had no analgesic effect in TASK-3 KO mice in all the tests. Collectively, our data suggest that the on-target activity of CHET3 is linked to the analgesic phenotypes.

Although CHET3 has a higher activation efficacy on TASK-3 over TASK-3/TASK-1, we suggest that both TASK-3 homomer and TASK-3/TASK-1 heteromer channels likely contribute to CHET3-induced analgesia for the following reasons: 1) *Kcnk9* is highly colocalized with *Kcnk3* in DRG; 2) TASK-3/TASK-1 heteromer has been found assembled efficiently and functionally in cerebellar granule cells (*45*), motoneurons (*46*) and carotid body glomus cells (*47*).

We found that CHET3 decreased the excitability without changing the RMP of nociceptive neurons. The lack of change in RMP could be explained by the fact that CHET3- or PK-THPP-mediated currents are negligible at approximately –60 mV. One may argue that there may be strong depolarizing “off-target” activity of CHET3 through another unknown channel/receptor, thereby masking the hyperpolarizing effect mediated by CHET3 on TASK-3-containing channels. However, if this were the case, one would at least expect PK-THPP to depolarize the RMP since PK-THPP, a molecule that is structurally distinct from CHET3, is unlikely to produce hyperpolarizing “off-target” activity through the same unknown channel/receptor.

CHET3 acted mainly on peripheral TASK-3-containing channels. Peripheral targets are much less likely to produce central side effects, including dependence/addiction. Although the utility of CHET3 and its derivatives as preclinical candidate compounds requires further assessment with systematic nonclinical safety tests performed in GLP (Good Laboratory Practice) in rodents and other animals, it seems that the activation of peripheral TASK-3-containing channels does not produce obvious severe acute side effects on the cardiovascular system, where TASK-3-containing channels are also expressed. Interestingly, we found that TASK-3 was more highly expressed in TG than in DRG. Further studies are needed to evaluate the translational potentials of TASK-3 activation (TASK-3/TASK-1) in TG to treat chronic pain related to trigeminal neuralgia and migraine. Finally, although TASK-3 is expressed in human DRG (*9*) and variation in *KCNK9* is involved in breast pain in breast cancer patients (*11*), direct evidence for the functional involvement of TASK-3 in pain signaling in humans is lacking. Future functional studies on human tissues or studies with genetic screening of TASK-3-related mutations in humans would greatly aid in assessing the translational potential of TASK-3 for treating pain in humans.

## MATERIALS AND METHODS

### Study design

Structure-based drug design methods were used to perform initial virtual screening, and patch-clamp electrophysiology was mainly used to study the activity/mechanism of candidate compounds on TASK-3-containing channels. The analgesic effects of TASK-3 activators were then studied in acute and chronic pain models in mice and rats. Pharmacokinetic analysis was performed to assess how CHET3 was distributed. KO mice were used to confirm the on-target activity of CHET3. Finally, *in situ* hybridization with the RNAscope technique was used to map the distribution of TASK-3 in DRG and TG. The functional roles of TASK-3 were assessed by measuring how CHET3 and PK-THPP modulate K^+^ currents, action potential firings and sensitivity to thermal stimulation in nociceptive neurons.

Sample size and replicates: For single-cell based experiments, at least 5 cells per condition were tested. For in *vivo* studies in animals, 6-10 animals per condition were used. No power analysis was performed to determine the sample size.

### Homology modeling for the TASK-3 structure

A sequence alignment was generated by using the Clustal Omega server (*16*). Notably, the two pore domains and selectivity filter sequence motifs were highly conserved among the K2P channels, which were largely used to guide the alignment. Conserved residues E30 and W78 in TASK-3 helped to locate the position of the non-conserved cap domain.

### Virtual screening

Docking was performed by using Schrödinger Glide software (New York, NY, USA). Compounds were screened using the high-throughput virtual screening (HVS) module followed by the standard docking module SP in Glide. The Glide G-score was used to rank the results list. To allow for diversity of molecular structures, binding modes and drug-like properties, twelve hits were selected for the bioassay.

### Chemicals

PK-THPP was purchased from Axon Medchem. CHET3 purchased from commercial sources was used in the initial electrophysiological screening. Then, CHET3 was synthesized in the lab for the following studies in this paper. The synthesis routes and characterization of CHET3 and its derivatives CHET3-1 and CHET3-2 are outlined in the Supplementary Materials.

For electrophysiology, stock solutions of CHET3 and its derivatives (50 mM) were prepared in dimethyl sulfoxide (DMSO) and diluted in the extracellular solution before use.

For animal studies, CHET3 and PK-THPP were both dissolved in 10% DMSO, 5% Tween 80 and 85% saline; CHET3-1 was dissolved in 10% DMSO, 5% ethoxylated castor oil, 35% poly (ethylene glycol) and 50% corn oil; and CHET3-2 was dissolved in 14% DMSO, 5% Tween 80 and 81% saline. The solvents were used as vehicle controls.

### Detailed modeling of the CHET3-TASK-3 binding poses

Initially, the configuration of CHET3 was determined by the Ligprep module in Schrödinger Maestro and Gaussain09 (Gaussian, Inc). Detailed descriptions are displayed in the Supplementary Materials. The configuration of the tautomer with the lowest energy was adopted to generate multiple ring conformations. CHET3 conformations were docked to the TASK-3 channel model by standard Glide as described for the virtual screening. Two binding modes (G-score values at −8.3 and −7.9, separately) were obtained from docking. In the best pose (1^st^ model in fig. S5A), the guanidyl group in CHET3 establishes a hydrogen bond with the backbone NH of residue L232 in TM2, while the guanidyl group in the additional mode of binding (2^nd^ model in fig. S5B) faces towards the selectivity filter and interacts with hydroxyl group of T199. To identify the accurate binding mode of CHET3, two docking models were further studied using molecular dynamics (MD) simulations (see below).

### MD simulations

The TASK-3 model obtained from homology modeling and two binding models of CHET3-bound TASK-3 were used to build the models of *apo* TASK-3 and CHET3-bound TASK-3, respectively. Models were inserted in a POPC (1-palmitoyl-2-oleoyl-sn-glycero-3-phosphocholine) lipid bilayer to establish the CHET3-bound system and the *apo* system, respectively. MD simulations were carried out by using GROMACS 5.1.4 (*48*) with CHARMM36 parameters (*49*).

### Comparison of the binding of CHET3, CHET3-1 and CHET3-2

The RosettaLigand application (*26, 27*) was applied to dock CHET3, CHET3-1 and CHET3-2. The best binding mode obtained from MD simulations was adopted as the initial docking model. For each docking trial, the top 1000 models were sorted by total score, and the binding energy between three compounds and the channel was calculated. Additionally, *in silico* alanine scans were conducted by individually changing the residue to alanine without otherwise changing the conformation of the protein or ligands in Rosetta. To explore the distribution of binding interactions between compounds and proteins, the average energies of the top 10 models with the lowest binding energies (interface score) were calculated. To compare the binding of CHET3, CHET3-1 and CHET3-2, the top 50 models of each compound with the lowest binding energies were used to calculate the total score and interface score.

### Electrophysiology

Electrophysiology tests of hTASK-3, hTASK-1, hTREK-1, mTREK-2, hTRAAK, hTHIK-1, hTRESK, hTASK-3/hTASK-1, hTRPM8 and hTRPV1 were performed with transiently transfected HEK-293T cells. The cDNAs of hTASK-3, hTASK-1, hTHIK-1, hTRESK and hTASK-3/hTASK-1 were subcloned into the pCDNA3 vector (Invitrogen). The cDNAs of hTREK-1, mTREK-2, hTRAAK, hTRPM8 and hTRPV1 were subcloned into the pEGFPN1 expression vector (Invitrogen). For hTASK-3/hTASK-1, concatemer products were designed for the 3’ and 5’ ends of TASK-3 and TASK-1, ensuring that the stop codon of TASK-3 was removed.

Electrophysiological tests of hERG, Kv2.1 and BK were performed with stable cell lines. The CHO-hERG stable cell line was generated in-house and was based on a standard CHO-K1 cell line. The HEK293-human Kv2.1 stable cell line and the CHO-human BK stable cell line were generated by Ion Channel Explore (Beijing, China). Whole-cell recordings of ion channels were performed with patch-clamp amplifiers (EPC10, HEKA or Axon 700B, Molecular Devices) at 23-25 °C. The current signals were filtered at 2 kHz and digitized at 10 kHz. The pipettes for whole-cell recordings were pulled from borosilicate glass capillaries (World Precision Instruments) and had a resistance of 3-7 MΩ. For recordings of K^+^ channels, the standard pipette solution contained (in mM) 140 KCl, 2 MgCl_2_, 10 EGTA, 1 CaCl_2_, and 10 HEPES (pH 7.3, adjusted with KOH), and the external solution contained (in mM) 150 NaCl, 5 KCl, 0.5 CaCl_2_, 1.2 MgCl_2_, and 10 HEPES (pH 7.3, adjusted with NaOH). For recordings of the TRPV1 and TRPM8 currents, the internal solution contained (in mM) 140 CsCl, 0.1 CaCl_2_, 1 MgCl_2_, 10 HEPES, and 5 EGTA (pH 7.2, adjusted with CsOH), and the external solution contained (in mM) 140 NaCl, 5 KCl, 1 MgCl_2_, 0.5 EGTA, and 10 HEPES (pH 7.4, adjusted with NaOH). For recordings of hERG, the outward current of hERG channels was elicited by a 2.5-second depolarization step to +30 mV from a holding potential of −80 mV, followed by a 4-second repolarization step to −50 mV to measure the tail current. For recordings of Kv2.1, currents were evoked by a 200-millisecond depolarization step to +60 mV from a holding potential of −80 mV. For recordings of BK, currents were evoked by a 1-second depolarization step to +70 mV from a holding potential of −80 mV. For recordings of TRPV1 and TRPM8, currents were recorded using a ramp protocol from −100 mV to +100 mV over a period of 400 milliseconds at a holding potential of 0 mV.

Single-channel current recording was acquired in the excised inside-out configuration of patch clamp using EPC10 (HEKA) at 23-25 °C. The pipettes had resistances of 7-15 MΩ. The standard pipette and bath solutions contained (in mM) 140 KCl, 1 CaCl2, 2 MgCl2, 10 HEPES and 10 EGTA (pH 7.4, adjusted with KOH). At acquisition the single-channel currents were low-pass filtered at 2 kHz and sampled at 10 kHz. Recordings lasting at least 50 s were put to further analysis to ensure enough events detected. A threshold at half the open channel current amplitude of the major conductance state was set to detect the single channel events. No junction potential correction was done. All the events in the selected section were detected automatically using Clampfit 10 (Molecular Devices, Inc.) followed by manual inspection. The amplitude histograms were fitted with Gaussian distributions with a bin width of 0.1 pA. TASK3 channel activity in an inside-out patch was expressed quantitatively as NP_O_ (N is the number of channels in the patch, and PO is the probability of a channel being open). The NP_O_ was calculated by the relative area under all points amplitude histogram and expressed as follows: NP_O_ = (A_1_ + 2A_2_ + 3A_3_ +…+ nA_n_) / (A_0_ + A_1_ + A_2_ +…+ A_n_), where A_0_ is the area under the Gaussian curve of an all points histogram corresponding to the closed state, A_1_…A_n_ represents the histogram area that indicates the level of the distinct open state for 1 to n channels in the patch examined, and n is the number of active channels. The single channel conductance of TASK3 channels was calculated using the ratio of current amplitude of the first open state to voltage at −60 mV or +60 mV.

### Ethics statement

All experiments with animals were approved by the Animal Research Committee of East China Normal University (PROTOCOL No. m20171020 and m20180112) and the Animal Research Committee of West China Hospital of Sichuan University (PROTOCOL No. 2018175A). For tissue collection, mice were given a lethal dose of pentobarbital intraperitoneally.

### Animals

BALB/c mice and Sprague-Dawley rats were used in most animal studies, and TASK-3 KO mice and WT control littermates were on a C57BL/6 background. Male mice or rats aged 8-10 weeks were used for behavioral tests unless stated otherwise. Animals were housed in a conventional facility at 21 °C on a 12 h light-dark cycle with unrestricted access to food and water.

### TASK-3 KO mice generation

To generate a *Kcnk9* knockout C57BL/6 mouse line with the CRISPR-Cas9 genome editing system, two single-guide RNAs (sgRNA-1, 5′-CCGCTTCATGGCCGCGAAGA<u>AGG</u>-3′, and sgRNA-2, 5′-AGGAACCGGCGAATTTCCAC<u>TGG</u>-3′) flanking exon1 were designed (Bioray Laboratories). A 241-bp deletion was bound to exon 1 of the *Kcnk9* gene locus, resulting in *Kcnk9*^Δ/Δ^ mice with a frameshift mutation. Additional information will be provided upon request.

### Spared nerve injury model

Unilateral spared nerve injury (SNI) surgery was performed. The experimental animals were placed in the prone position. After disinfection with povidone iodide and 75% ethanol, a minimal skin incision was made at the mid-thigh level to expose the sciatic nerve and its three branches by separating the muscle layers. The tibial and common peroneal nerves were tightly ligated with 5.0 silk threads, and a 1-2 mm section was removed between the proximal and distal parts of the nerves. The sural nerve was restrictively preserved to avoid any harmful injury. The muscle layer and skin were closed after surgery, and the animals were transferred to a warm pad to recover from anesthesia.

### Chronic inflammatory pain model

A volume of 20 μL Complete Freund’s Adjuvant (CFA) (Sigma-Aldrich) was subcutaneously injected into the left hindpaw of the mice to induce chronic inflammatory pain in mice. After injection, the syringe was maintained for at least 30 s to avoid overflow.

### Tail immersion

Mice were restrained in the test tube with their tails stretching out and moving freely 15 min twice daily for 3 days. The distal third of the tail was immersed in a water bath at 5 °C, 20 °C, 40 °C, 46 °C, or 52 °C. Three measurements of tail flick latency (in seconds) to stimulation, as indicated by rapid tail flexion, were averaged. A cutoff value of 15 seconds was adopted to prevent unexpected damage.

### Formalin test

Mice were housed individually in Plexiglas chambers. After habituation to the testing environment for at least 30 min, the left hindpaw of the mice was injected subcutaneously with formalin (20 μL of 2.5% formalin, diluted in saline), and the mice were placed into the chamber of the automated formalin apparatus, where movement of the formalin-injected paw was recorded by an action camera (SONY, HDR-AS50). The number of paw flinches was counted at 5 min intervals for 60 min by a blind experimenter. The time spent exhibiting these pain behaviors was recorded for the first (0-10 min) and second phases (10-60 min).

### Paw pressure

The effects of mechanical nociception were evaluated with an Analgesimeter (model 37215; Ugo-Basile, Varese, Italy). Mice were placed in the testing room for 3 continuous days to acclimate to the environment. The hindpaws of mice were pressed with a constant pressure of 450 g using a cone-shaped paw-presser with a rounded tip and immediately released as soon as the animal showed a struggle response, and the reaction latency was recorded in seconds. The analgesic effects of TASK-3 agonists were evaluated 30 min after i.p. injection.

### Spontaneous pain test

After 3 days of acclimation, the SNI mice were placed in an elevated transparent cage (20 × 20 × 14 cm) with a wire mesh floor (0.5 × 0.5 cm). A 5 min duration was videoed by an action camera (SONY, HDR-AS50) for each mouse, and the number of left hindpaw flinches was calculated by a blind experimenter.

### Cold plantar test

Mice were allowed to acclimate to the testing environment 2-3 h daily for 3 continuous days. The cold probe produced freshly with fine dry ice powder in a 5 mL syringe was held against a 6 mm depth of flat glass. The center of the hindpaw was targeted, and the withdrawal latency, manifesting as a quick flick or paw licking, was recorded. A cutoff time of 30 s was used to prevent potential tissue damage.

### von Frey test

The SNI rats and mice were individually placed in the chamber as described in the spontaneous pain test. Mechanical sensitivity was assessed by two methods.

Method 1 (for all the von Frey tests described except Fig. 5D): The mechanical paw withdrawal threshold was assessed using von Frey filaments with an ascending order. The tip of the filament was perpendicularly targeted to the region comprising the sural nerve territory, and sufficient stimulation was maintained for 1 s. Rapid paw withdrawal or flinching was considered a positive response, and the bending force for which at least 60% of the application elicited a positive response was recorded as the mechanical paw withdrawal threshold.

Method 2 (up-down method): Mechanical responses were tested by stimulating the region comprising the sural nerve territory with von Frey monofilaments by using the up-and-down method, starting with 0.04 g. Biting, licking, and withdrawal during or immediately following the 3 s stimulus were considered a positive response.

### Hargreaves test

Hindpaw sensitivity to a noxious thermal stimulus was assessed using a radiant heat source (model 37370; Ugo-Basile, Varese, Italy). The stimulus intensity was set to produce an approximate latency of 10 s at baseline, and a cut-off value was set at 20 s to avoid unexpected damage. Mice were allowed to acclimate in Plexiglas chambers with a glass floor for 3 days, and the time to paw withdrawal was measured per mouse with a 5 min inter-stimulation period. Three trials were averaged to yield the withdrawal latency.

### RNAscope *in situ* hybridization

The sequences of the target probes, preamplifier, amplifier and label probes are proprietary and commercially available (Advanced Cell Diagnostics). *In situ* hybridization was performed on frozen DRG sections (10 μm) using RNAscope Multiplex Fluorescent Reagent Kit v2 (ACDbio, Cat#323100) and RNAscope 4-Plex Ancillary Kit for Multiplex Fluorescent Kit v2 (ACDbio, Cat#323120). The hybridization assay was performed as described by the vendor^’^s protocol. The *in situ* probes included *Kcnk3* (Cat#534881), *Kcnk9* (Cat#475681), *Trpa1* (Cat#400211), *Trpv1* (Cat#313331), *Trpm8* (Cat#420451), *Rbfox3* (Cat#313311), *Th* (Cat#317621), *Ntrk2* (Cat#423611), and *P2rx3* (Cat#521611). The specificity of the fluorescence signals was validated by an RNAscope 3-plex Positive Control Probe (Cat#320881) and an RNAscope 3-plex Negative Control Probe (Cat#320871). Fluorescence images were taken using a NIKON A1R^+^MP two-photon confocal scanning microscope and were analyzed using ImageJ software.

### Acutely dissociated DRG neuron preparation and electrophysiology

Three to six-week-old male Sprague-Dawley rats were sacrificed. The DRGs were collected in a 35-mm tissue culture dish and digested in 3% collagenase for 20 min, followed by 1% trypsin for another 30 min. After titrating by sucking up and down, the DRG neurons were cultured in neurobasal medium containing 2% B27 medium for 2-4 h. The bath solution contained (in mM) 140 NaCl, 3 KCl, 1.3 MgCl2, 10 HEPES, 2.4 CaCl2, and 10 glucose, pH 7.3. The pipette solution contained (in mM) 40 KCl, 10 HEPES, 5 EGTA, 10 NaCl, 95 K-gluconate, and 4 Mg-ATP, pH 7.4. To minimize voltage-gated currents, voltage ramps from −120 mV to −30 mV were applied, and 1 mM CsCl was added extracellularly to block hyperpolarization-activated currents. To determine the reversal potential of the CHET3-sensitive currents, NaCl was replaced with equimolar KCl. Whole-cell recordings were performed with hardware settings similar to those described for electrophysiology in HEK-293T cells.

### Acutely dissociated DRG neuron preparation and intracellular Ca^2+^ imaging

Two-week-old male Sprague-Dawley rats were sacrificed. The DRGs were collected in a 35-mm tissue culture dish and digested in 2.5 mg/mL papain (Sigma-Aldrich) for 30 min at 37 °C, followed by 2.5 mg/mL collagenase (Sigma-Aldrich) for another 30 min. Digested ganglia were gently washed with neurobasal medium and mechanically dissociated by passage through the pipette. Neurons were seeded on laminin-coated wells (Corning) and cultured overnight at 37 °C in 5% CO_2_ in neurobasal medium supplemented with 2% B27 (Sigma-Aldrich) and containing 50 ng/mL GDNF (PeproTech) and 50 ng/mL BDNF (PeproTech).

Changes in intracellular Ca^2+^ concentration were monitored using ratiometric Fura-2-based fluorimetry. Neurons were loaded with 2 μM Fura-2-acetoxymethyl ester (Yeasen) dissolved in bath solution and 0.02% Pluronic F127 (Sigma-Aldrich) for 30 min at 37°C. Fluorescence was measured during alternating illumination at 340 and 380 nm using an Olympus IX73 inverted fluorescence microscopy system. The bath solution contained (in mM) 138 NaCl, 5.4 KCl, 2 CaCl2, 2 MgCl2, 10 glucose, and 10 HEPES, pH 7.4, adjusted with NaOH. At the end of each experiment, cells were subjected to a depolarizing solution containing 50 mM KCl, and cells nonresponsive to 50 mM KCl were excluded from the analysis. The bar graphs in Fig. 7I and fig. S15B are pooled data from both responding cells and nonresponding cells under different conditions.

### Thermal stimulation

Coverslip pieces with cultured cells were placed in a recording chamber and continuously perfused (approximately 1 mL/min).

Cool stimulation: The temperature was adjusted with iced perfusion solution and controlled by a feedback device. Cold sensitivity was investigated with an ∼2 min duration ramp-like temperature drop from 37 °C to ∼15 °C.

Heat stimulation: The temperature was adjusted with a water-heated device (model TC-324B/344B, America), with the temperature of the perfusion solution raised and controlled by a feedback device. Heat sensitivity was investigated with an ∼5 min duration ramp-like temperature rise from 25 °C to ∼43 °C.

### Statistical analysis

Statistical analyses were carried out using Origin 9.0 software (Origin Lab Corporation, Northampton, USA). Data were analyzed as described in the figure legends. The normality of the data distribution was determined before appropriate statistical methods were chosen. The drug was assessed as significantly active by using statistical tests to compare the values at baseline to those at given time points unless specified. No statistical methods were used to predetermine sample sizes.

## SUPPLEMENTARY MATERIALS

### Materials and Methods

Fig. S1. A potential druggable pocket identified in several structures of K2P channels.

Fig. S2. Selectivity of PK-THPP against hERG, Kv2.1 and BK channels.

Fig. S3. Representative current traces for whole-cell recordings on several K2P channels and other ion channels.

Fig. S4. 10 μM CHET3 does not show effect on three pain-related GPCRs.

Fig. S5. Binding modes of CHET3 suggested by docking and MD simulations.

Fig. S6. Whole-cell path-clamp current recording for three TASK-3 mutants.

Fig. S7. Conformations of the extracellular ion pathway in MD simulations.

Fig. S8. Dose-dependent analgesia by CHET3 in mechanical allodynia.

Fig. S9. Effects of CHET3 on the locomotion activities, blood pressure and body temperature in rodents.

Fig. S10. Comparison of the binding of CHET3, CHET3-1 and CHET3-2.

Fig. S11. Blockade of CHET3-1 analgesia by PK-THPP.

Fig. S12. Generation and characterization of TASK-3 gene (*Kcnk9*) knockout mice.

Fig. S13. Down-regulation of peripheral TASK-3 under chronic pain.

Fig. S14. Effects of CHET3 and PK-THPP on nociceptive neurons.

Fig. S15. Thermal stimulation induced Ca^2+^ signals were mediated by TRP channels.

Table S1. Echocardiographic evaluation of CHET3 on mice.

Table S2. CHET3 pharmacokinetics in plasma and brain following a single intraperitoneal administration to naïve male C57BL/6 mice.

Table S3. CHET3 pharmacokinetics in plasma and brain following a single intraperitoneal administration to SNI 7-d male C57BL/6 mice.

## Acknowledgements

We thank Bioray Laboratories for technical support in preparing *Kcnk9* knockout mice. We thank Dr. Tao Li (West China Hospital) for technical assistance with blood pressure test and Miss Rongrong Cui (SIMM) for assistance on PK test. We thank the supports of ECNU Multifunctional Platform for Innovation (001 and 011).

## Author contributions

P.L. performed behavior tests, ISH, Ca^2+^ imaging; Y.Q. performed drug design and computations; Y.M. performed mouse genetics on TASK-3 KO mice and behavior tests; J.F. performed electrophysiology; Z.S. performed electrophysiology, behavior tests, ISH; L.H. performed electrophysiology; S.B., Y.W. and B.S. performed Ca^2+^ imaging; J.Z. and W.G.L. performed elevated plus maze tests; Z.C. and N.P. assisted with behavior tests and cell culture; E.S. performed dark/light box tests; L.Y. assisted with behavior tests; F.T., X.L. and Z.G. performed electrophysiology for some of the initial compound screenings; P.S., Y.C. and Y.M. performed pharmacokinetics study; D.H. performed the qPCR experiments for TASK-3 KO mice; L.Z. performed experiments of μOR; D.Y. performed experiments of 5-HT_1B_R; W.L. performed experiments of CB1R; T.Y., J.X. and Y.M. performed experiments of echocardiography. Q.Z. prepared the derivatives of CHET3. J.L. oversaw the animal behavior tests. H.J. oversaw the computations. R.J. and H.Y. initiated, supervised the project, analyzed the experiments, and wrote the manuscript with input from all co-authors.

## Data and materials availability

All data is available in the main text or the supplementary materials.

