## Supplementary material for "Selective Activation of TASK-3-containing K^+^ Channels Reveals Their Therapeutic Potentials in Analgesia": Methods, Supplemental Figures and Tables

**Supplementary Materials includes:**

Materials and Methods

Fig. S1. A potential druggable pocket identified in several structures of K2P channels.

Fig. S2. Selectivity of PK-THPP against hERG, Kv2.1 and BK channels.

Fig. S3. Representative current traces for whole-cell recordings on several K2P channels and other ion channels.

Fig. S4. 10 μM CHET3 does not show effect on three pain-related GPCRs.

Fig. S5. Binding modes of CHET3 suggested by docking and MD simulations.

Fig. S6. Whole-cell path-clamp current recording for three TASK-3 mutants.

Fig. S7. Conformations of the extracellular ion pathway in MD simulations.

Fig. S8. Dose-dependent analgesia by CHET3 in mechanical allodynia.

Fig. S9. Effects of CHET3 on the locomotion activities, blood pressure and body temperature in rodents.

Fig. S10. Comparison of the binding of CHET3, CHET3-1 and CHET3-2.

Fig. S11. Blockade of CHET3-1 analgesia by PK-THPP.

Fig. S12. Generation and characterization of TASK-3 gene (*Kcnk9*) knockout mice.

Fig. S13. Down-regulation of peripheral TASK-3 under chronic pain.

Fig. S14. Effects of CHET3 and PK-THPP on nociceptive neurons.

Fig. S15. Thermal stimulation induced $Ca^{2+}$ signals were mediated by TRP channels.

Table S1. Echocardiographic evaluation of CHET3 on mice.

Table S2. CHET3 pharmacokinetics in plasma and brain following a single intraperitoneal administration to naïve male C57BL/6 mice.

Table S3. CHET3 pharmacokinetics in plasma and brain following a single intraperitoneal administration to SNI 7-d male C57BL/6 mice.

### MATERIALS AND METHODS

#### Protonation states of CHET3

Initially, given that various conjugated tautomers exist in guanidyl-pyrimidine moiety, correct tautomeric states and protonation states of CHET3 were generated at pH 6-8 by Ligprep module in the Schrödinger Maestro. Two tautomers were generated and presented as follows:

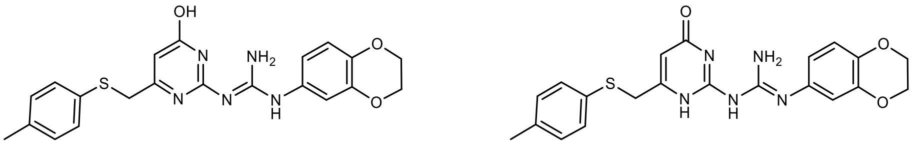


State 1 State 2

The energy of each state was calculated using Gaussain09 (Gaussian, Inc) at RB3LYP/def2tzvp. The energy of State 1 was 0.377 kJ/mol lower than that of State 2. This result that State 1 has a lower energy than State 2 is also consistent with a previous report of electronic and structural analysis of biguanide derivatives (*50*). Thus, State 1 was adopted in the current work.

#### Synthesis of CHET3 ((*E*)-1-(2,3-dihydrobenzo[*b*][1,4]dioxin-6-yl)-2-(4-hydroxy-6-((*p*-tolylthio)methyl)pyrimidin-2-yl)guanidine)

Scheme：

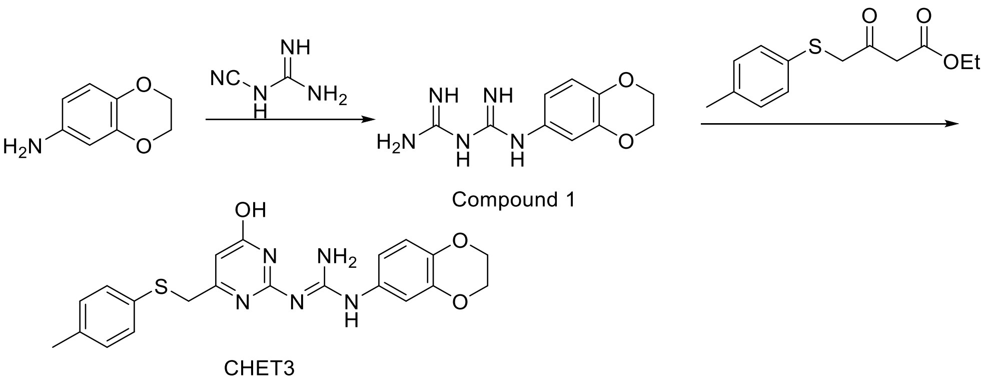


Step 1. 1-(2,3-dihydrobenzo[*b*][1,4]dioxin-6-amine)biguanide (Compound 1). The mixture of 2,3-dihydrobenzo[b][1,4]dioxin-6-amine (3.02 g, 20 mmol) , dicyandiamide (1.68 g, 20 mmol) and 37% aq HCl (2 mL) in 1,4-dioxane (30 mL) was stirred at reflux for 3 hours. TLC showed the reaction was completed. The reaction mixture was cooled to room temperature and added saturated $NaHCO_3$ to pH 6-7. The resulting precipitated solid was filtrated at this temperature, the solid was washed with EtOH, dried to obtain the desired product Compound 1. Yield: 2.05 g (44%). LCMS: 236.1$[M+1]^+$.

Step 2. (*E*)-1-(2,3-dihydrobenzo[*b*][1,4]dioxin-6-yl)-2-(4-hydroxy-6-((*p*-tolylthio)methyl)pyrimidin-2-yl)guanidine (CHET3). The mixture of Compound 1 (700 mg, 2.97 mmol) and ethyl 3-oxo-4-(*p*-tolylthio)butanoate (1.21 g, 4.8 mmol) in anhydrous EtOH (20 mL) was heated to reflux for 6 hours. The reaction mixture was cooled to room temperature and the precipitated solid was collected by filtration. The crude product was purified by pre-HPLC to obtain the desired product CHET3. Yield: 750 mg (60%). LC-MS: 424.1$[M+1]^+$, Purity: 99.8%; $^1$HNMR (400 MHz, *DMSO_d6*): δ 11.20 (br, 1H), 9.25 (s, 1H), 7.23(d, J = 8.0Hz, 2H), 7.09(d, J = 8.0Hz, 2H), 7.01 (d , J = 2.0Hz, 1H), 6.95(d, J =

8.8Hz, 1H), 6.74 (d, J = 8.4Hz, 1H), 5.64 (s, 1H), 4.19 (q, J = 2.8 Hz, 4H), 3.88 (s, 2H), 2.23 (s, 3H). $^{13}$C NMR (400 MHz, *DMSO_d6*): δ 164.4 (C), 162.7 (C), 159.1 (C), 156.9 (C), 143.5 (C), 139.9 (C), 136.0 (C), 132.4 (CH), 130.1 (CH), 129.4 (CH), 125.1 (CH), 117.2 (CH), 115.1 (CH), 110.8 (CH), 104.0 (CH), 64.5 ($CH_2$), 64.4 ($CH_2$), 43.6 ($CH_2$), 21.0 ($CH_3$). HRMS-ESI (m/z) $[M + H]^+$ Calculated for for $C_{21}H_{21}N_5O_3S$: 423.1465 found: 424.1442.

**Synthesis of CHET3-1 ((*E*)-2-(4-hydroxy-6-((*p*-tolylthio)methyl)pyrimidin-2-yl)-1-(naphthalen-1-yl)guanidine)**

Scheme:

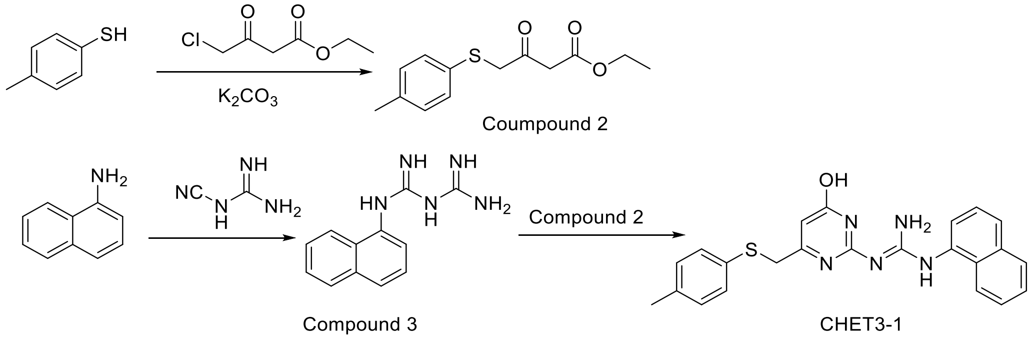


Step 1. ethyl 3-oxo-4-(*p*-tolylthio)butanoate (Compound 2). To a solution of 4-methylbenzenethiol (4.26 g, 34.3 mmol) in acetone (50 mL), $K_2CO_3$ (12.0 g, 86.8 mmol) and ethyl 4-chloro-3-oxobutanoate (4.70 g, 28.6 mmol) were added at room temperature. The resulting mixture was stirred at room temperature for 3 hours. TLC showed the reaction was completed. The reaction mixture was poured into ice water, extracted with EtOAc, the combined organic phase was washed with water, brine, dried over anhydrous $Na_2SO_4$, concentrated. The crude product was purified with silica gel chromatograph to obtain the

desired product Compound 2 as a light yellow oil. Yield: 6.52 g (90%). LCMS: 253.1[M+1]$^+$; $^1$HNMR: (CDCl3, 400MHz): 1.27 (t, J= 6.8Hz, 3H), 2.32(s, 3H), 3.63 (s, 2H), 3.75(s, 2H),4.18(q, J= 6.8Hz, 2H),7.12(d, J =8.0Hz,2H), 7.27(d, J =8.0Hz,2H).

Step 2. 1-(1-naphthalen)-biguanide (Compound 3). The mixture of naphthalen-1-amine (1.43 g, 10 mmol), dicyandiamide (0.84 g, 10 mmol), 37% aq HCl (1 mL) in 1,4-dioxane (20 mL) was heated to reflux for 16 hours. TLC showed the reaction was completed. The mixture was cooled to room temperature, and the precipitated solid was collected by filtration. The solid was dissolved in saturated $NaHCO_3$ and the resulting solid was filtrated, washed with water, dried to obtain the desired product Compound 3. Yield: 1.86 g (82%) as a yellow solid. LCMS: 228.2[M+1]$^+$; $^1$HNMR (400 MHz, *DMSO_d6*): δ 8.03 (d, J =8.0 Hz, 1H), 7.83 (d, J = 8.0 Hz, 1H), 7.51 – 7.33 (m, 4H), 7.04 (br, 1H).

Step 3. (*E*)-2-(4-hydroxy-6-((*p*-tolylthio)methyl)pyrimidin-2-yl)-1-(naphthalen-1-yl) guanidine (CHET3-1). The mixture of Compound 2 (1.60 g, 6.34 mmol), Compound 3 (1.00 g, 4.40 mmol) in anhydrous EtOH (10 mL) was heated to reflux for overnight. The resulting precipitated solid was filtrated at this temperature, and the solid was washed with EtOH, dried to obtain the desired product CHET3-1. Yield: 820 mg (45%). LC-MS: 416.2[M+1]$^+$, Purity: 99.6%; $^1$HNMR: (400 MHz, *DMSO_d6*): δ 2.20 (s, 3H), 3.89(s, 2H), 5.65(s, 1H), 7.04(d, J =8.0Hz, 2H),7.19(d, J = 8.0Hz, 2H),7.49-7.59(m, 3H), 7.74(br,1H), 7.80 (d, J = 8.4Hz, 1H), 7.95-7.97(m, 1H),8.03-8.05(m, 1H),10.03(br, 1H),11.14(br, 1H).$^{13}$C NMR (400 MHz, *DMSO_d6*): δ 164.8 (C), 162.6 (C), 159.2 (C), 158.4 (C), 136.0

(C), 134.4 (C), 132.3 (C), 130.0 (CH), 129.4 (CH), 129.0 (CH), 128.7 (CH), 126.8 (CH), 126.7 (CH), 126.5 (CH), 126.3 (CH), 125.2 (CH), 123.5 (CH), 122.8 (CH), 104.0 (CH), 43.6 ($CH_2$), 21.0 ($CH_3$). HRMS-ESI (m/z) $[M + H]^+$ Calculated for $C_{23}H_{21}N_5OS$: 416.1467 found: 416.1548.

**Synthesis of CHET3-2 ((*E*)-1-(4-(tert-butyl)phenyl)-2-(4-hydroxy-6-((*p*-tolylthio)methyl)pyrimidin-2-yl)guanidine)**

Scheme：

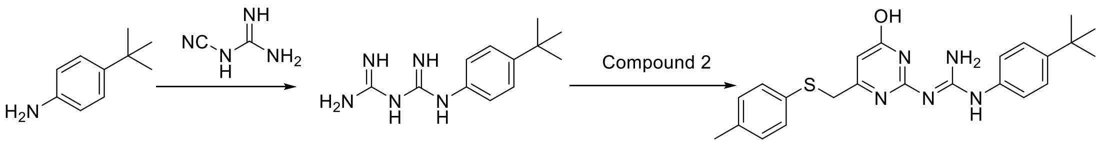


Compound 4 CHET3-2

Step 1. 1-(4-(tert-butyl)aniline)-biguanide HCl salt (Compound 4). The mixture of 4-(tert-butyl)aniline (3.00 g, 20.1 mmol) and dicyandiamide (1.68 g, 20 mmol), 37% aq HCl (2 mL) in 1,4-dioxane (20 mL) was heated to reflux for 10 hours. TLC showed the reaction was completed. The reaction mixture was concentrated to get the crude product Compound 4, which was used to next step without further purification. Yield: 3.12g (crude). LCMS： 234.1$[M+1]^+$.

Step 2. (*E*)-1-(4-(tert-butyl)phenyl)-2-(4-hydroxy-6-((*p*-tolylthio)methyl)pyrimidin-2-yl)guanidine (CHET3-2). The mixture of Compound 2 (1.10 g, crude), Compound 4 (1.00 g, 4.0 mmol), $NaHCO_3$ (200 mg, 2.38 mmol) in anhydrous EtOH (30 mL) was heated to reflux overnight. The reaction mixture was cooled to room temperature and the precipitated solid was cooled by filtration. The crude product was purified by pre-HPLC to obtain the desired product CHET3-2. Yield: 416 mg (25%). LC-MS: 422.2$[M+1]^+$, Purity: 99.7%; $^1$HNMR (400 MHz, *DMSO_d6*): δ 11.15 (s, 1H , 9.06 (s, 1H , 7.52(d, J = 8.4Hz, 2H), 7.27(t, J = 8.0Hz, 4H), 7.12(d, J = 8.0Hz, 2H), 5.67(s, 1H), 3.93(s, 2H), 2.26(s, 3H), 1.27(s, 9H). $^{13}$C NMR (400 MHz, *DMSO_d6*): δ 164.3 (C), 162.7 (C), 159.1 (C), 156.7 (C), 145.5 (C), 136.5 (C), 136.0 (CH), 132.4 (CH), 130.1 (CH), 129.4 (CH), 125.8 (CH), 125.1 (CH), 121.0 (CH), 104.1 (CH), 43.6 ($CH_2$), 34.4 (C), 31.7 ($CH_3$), 21.0 ($CH_3$). HRMS-ESI (m/z) $[M + H]^+$ Calculated for $C_{23}H_{27}N_5OS$: 422.1936, found: 422.2004.

**cAMP assay of human μOR**

HEK293 cells were cultured for forty-eight hours in 6-well plates followed by transient transfection of pCDNA3-HA-OPRM1 plasmid. Then the cells were incubated with non-enzyme cell dissociation solution (Sigma) at 37 °C for 10 min. After centrifugation, the cells were suspended in DMEM containing 500 μM 3-isobutyl-1-methylxanthine (IBMX, Sigma). Then, the cells were transferred into a 384-well plate at a density of 10000 cells per well. The cells were treated with Forskolin (1.25 μM) and/or compounds at room temperature for 30 min. μOR agonist DAMGO (MedChemExpress) was used as a reference. The cells were incubated with cAMP-$D_2$ and anti-cAMP Crypate (cAMP kit, Cisbio) for 1 h at room temperature and readout using PerkinElmer Envision.

**cAMP assay of human 5-$HT_{1B}$R**

CHO-5-$HT_{1B}$R stable cell line was established in-house based on a standard CHO-K1 cell line. Briefly, cells stably expressing 5-$HT_{1B}$R were seeded into white poly-D-lysine coated

384-well plates at a density of 2,000 cells per well for 24 h at 37 °C. The cells were treated with Forskolin (1 μM) and/or compounds at room temperature for 30 min. 5-$HT_{1B}R$ agonist dihydroergotamine (DHE, Tocris) was used as a reference. The cells were incubated with cAMP-$D_2$ and anti-cAMP Crypate (cAMP kit, Cisbio) for 1 h at room temperature and readout using PerkinElmer Envision.

**Calcium flux assay of human $CB_1R$**

CHO cells expressing $CB_1R$ and Gα16（25,000 cells/100 μL）were seeded into 96-well plates (Corning, Painted Post, NY) and then incubated with Calcium-5 assay kit (Molecular Devices, Silicon Valley, CA). Compounds were injected using the Flexstation III instrument (Molecular Devices, Silicon Valley, CA). $CB_1R$ agonist WIN 55,212-2 (MedChemExpress) was used as a reference. Intracellular calcium flux was recorded for 120 s.

**Elevated plus maze test**

The EPM was made of grey acryl glass and elevated at a height of 50 cm above the floor. It consisted of four equally spaced arms (30 × 6 cm) radiating out from a central area (6 × 6 cm). Two opposing closed arms were enclosed from all sides except for the side adjoining the central area, and the remaining two open arms were exposed. A digital camera was mounted above the maze to record the images, which were quantified using the Ethovision video tracking system (Noldus Information Technology, Wageningen, Netherlands). Trials, which lasted 5 min, began once an animal was placed gently in the center area, facing one of the open arms. After each trial, the entire maze was cleaned and animal was returned to the home cage.

**Light/dark box test**

The light-dark apparatus consisted of a Plexiglas box (30 × 30 × 60 cm) separated into two compartments: a half for dark section and another for light section. Mice were initially placed in the center of the light section and facing away from the partition allowing them to move freely for 10 min. When all paws were positioned within the compartment, a mouse was considered to have entered. The number of light–dark transitions, time spent in the light or dark section, and latency to enter dark were recorded by a trained observer.

**Open field test**

We conducted the open field test in a square Plexiglas apparatus (40 × 40 × 35 cm) under diffused lighting. In detail, the arena was partitioned such that there was a “center” zone (20 × 20 cm) and a “corner” zone occupying the remaining area (*51*). A digital camera was set directly above the apparatus. Images were captured at a rate of 5 Hz and quantified using the Ethovision video tracking system (Noldus Information Technology, Wageningen, Netherlands). Mice were gently placed in the center of the square and allowed to explore freely for 5 min. After each trial, the apparatus was cleaned and the animal returned to the home cage.

**Rotarod test**

In the rotarod assay, mice were initially placed on the stationary rod. After habituation, rotation was started at 4 r.p.m (rotations per min), accelerating over a 5-min period to 40 r.p.m. The time taken for the mouse to fall from the rod was recorded.

**Grip strength test**

Forelimb strength was determined using a grip strength meter for mice (model 47200; Ugo Basile, Varese, Italy) according to previously described methods (*56*, *57*). Mice were accustomed to handling with leather gloves for three days before experiments were conducted, five minutes for each mouse per day. Mice were also accustomed to placement of forelimbs on the apparatus trapeze for several times before actual measurements were performed. For data acquisition, a mouse was suspended gently by the tail and lowered so that it could grasp the force transducer (trapeze) of a grip-strength meter for mice. The peak force in grams achieved at the time the animal lost its grip was stored and displayed by a peak preamplifier. Five trials were played with an interval of 3 min (for the mouse to rest and recover), and the results of middle three trials were analyzed.

**Rectal temperature test**

The test was carried out by inserting the thermistor probe of a digital thermometer (TH-212) to a depth of 1 cm into the rectum of the animals lubricated with petroleum jelly. The rectal temperature was recorded just before (0 min) the administration of test dose and at 30, 60, 90, and 120 min after administration of test doses. The ambient temperature was 23.2 ± 1 °C.

**Blood pressure test**

We performed the test using a non-invasive measurement system (Visitech BP-2000, USA) by tail-cuff method. The measurement was conducted on conscious rats closed in measuring tubes based on the measurement of the pulse in the tail artery. Rats were habituated to the touch of a human hand and to the measuring tubes one week before initiation of the experiment. For each rat, one measurement session comprised 5 readings of systolic and diastolic blood pressure and the arithmetic mean was calculated.

**Echocardiography**

The hair on the chest was removed with hair removal gel to minimize resistance to ultrasonic beam transmission. 45 min post dosing, transthoracic echocardiography was performed using the Vevo 2100 ultrasound system (Visualsonics, Toronto, Canada) equipped with a high-frequency (40 MHz) linear array transducer. Animals were placed supine on an electrical heating pad at 37°C under light isoflurane anesthesia (usual maintenance level 1.5% isoflurane / 98.5% oxygen). Continual ECG monitoring was obtained via limb electrodes. Heart rates were 450 ± 20 beats/min for the duration of the study (with adjustment of level of anesthesia as necessary). A two-dimensional echocardiographic and M-mode study was performed predominantly in parasternal long-axis. Analysis of LV volumes, ejection fraction (EF %) and fractional shortening (FS %) was performed with Vevo 2100 software (version 2.2.0) by one experienced cardiologists blinded to the treatment.

**Pharmacokinetics study**

Pharmacokinetics of CHET3 was analyzed in naive male C57BL/6 mice (n = 6) and male C57BL/6 mice 7 days after SNI (n=6). Plasma and brain concentrations were determined using LC-MS/MS methods after a single intraperitoneal injection dose (i.p. 10 mg/kg) of compound as a clear solution in 10% DMSO + 5% Tween80 + 85% saline at a concentration of 0.5 mg/mL. Blood samples were collected into heparinized test tube at each time point (0.083 h, 0.25 h, 0.5 h, 1.0 h, 2.0 h, 4.0 h and 8.0 h) and centrifuged at 2,000 g for 15 minutes to generate plasma samples. Brains were collected after myocardial perfusion with normal saline and homogenated with brain: normal saline (1:2, W/V) to generate brain samples. LC-MS/MS methods to quantify CHET3 in plasma and brain samples were developed. Plasma and brain samples were extracted with ACN, mixed and centrifuged. The supernatants were transferred into injection vials. The samples were analyzed with a Shimadzu HPLC system coupled with a 4000 Qtrap mass spectrometer (AB SCIEX, USA), which was equipped with an Applied Biosystems electrospray ionization (ESI) source and operated with Analyst 1.6.3 (AB SCIEX, USA). The column used was an Agela Venusil XBP C18 50 × 2.1 mm, 5 micron column. The mobile phase consisted of A: 5 mM $NH_4OAc$ with 0.1% formic acid in water, and B: MeOH. Standard curves were prepared by spiking compounds into control plasma and brain and these were used to determine drug concentrations. Pharmacokinetic parameters including area under the curve (AUC), $T_{max}$, $C_{max}$, and $T_{1/2}$ were calculated via non-compartmental analysis (NCA) using Phoenix WinNolin version 6.4 with mean concentration at each time point.

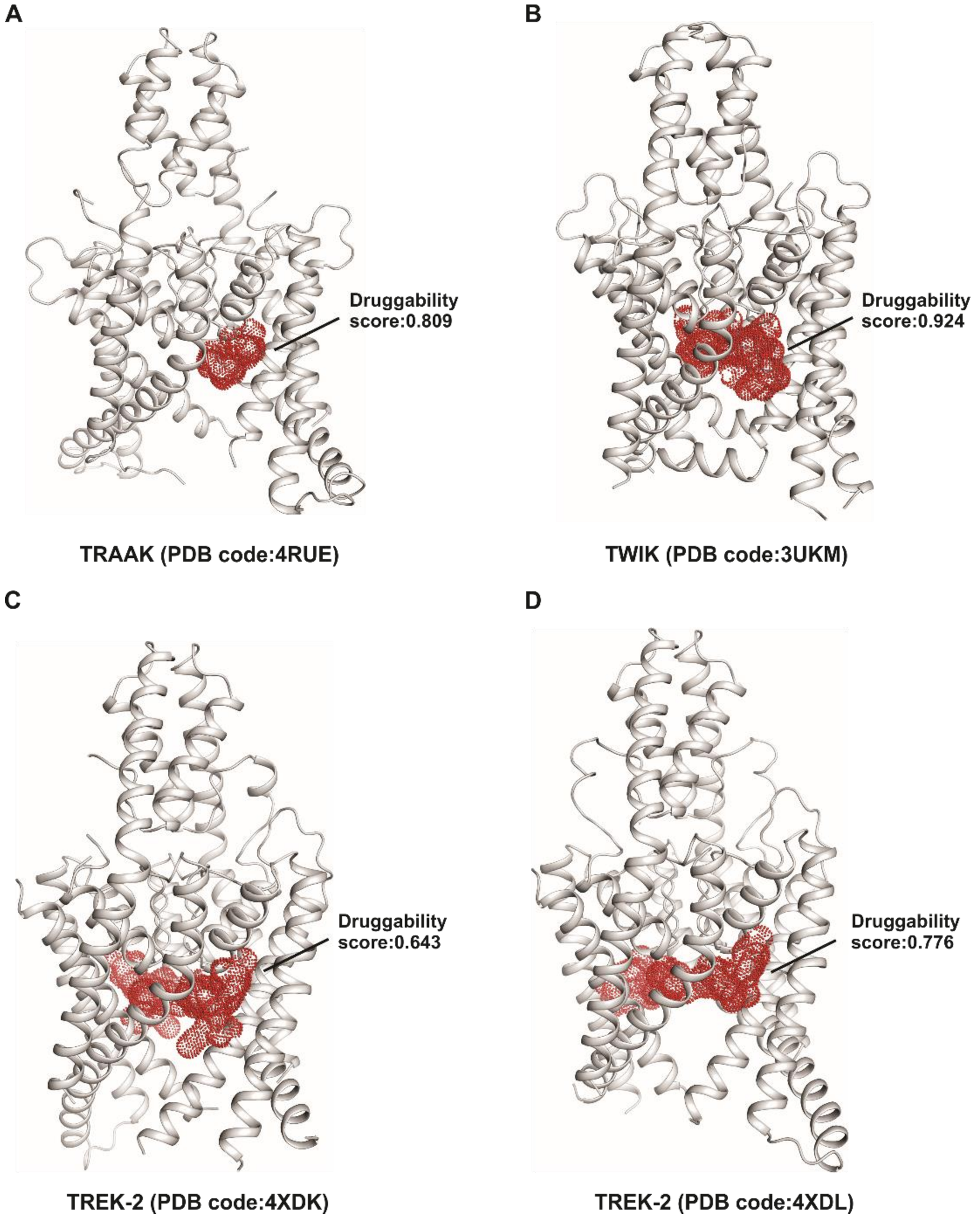


**Fig. S1. A potential druggable pocket identified in several structures of K2P channels.**

The pockets in four structures (PDB codes 4RUE (**A**), 3UKM (**B**), 4XDK (**C**) and 4XDL (**D**)) (*13*-*15*) were exhibited in red mesh. Druggability scores of the pockets were labeled.

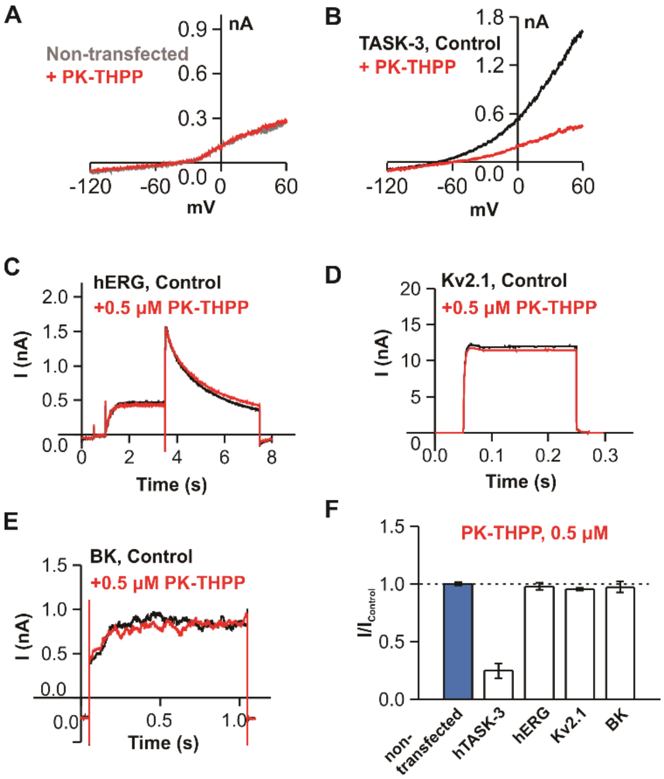


**Fig. S2. Selectivity of PK-THPP against hERG, Kv2.1 and BK channels.** (**A** to **E**) Representative current traces for TASK-3, hERG, Kv2.1 and BK channels in response to 0.5 µM PK-THPP. (**F**) Summary for the effects of PK-THPP on TASK-3, hERG, Kv2.1 and BK channels (n = 5-6).

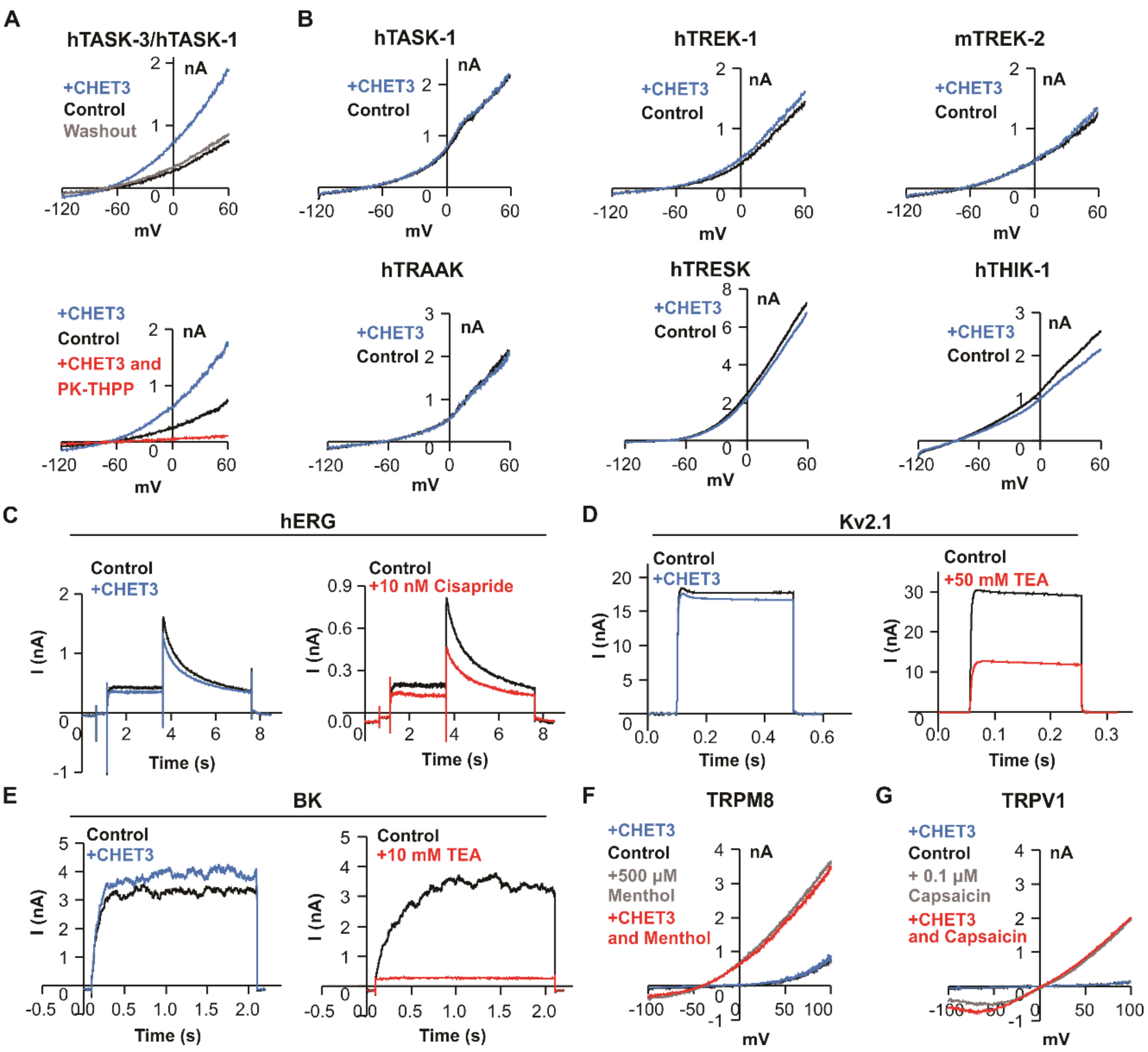


**Fig. S3. Representative current traces for whole-cell recordings on several K2P channels and other ion channels.** (**A**) Representative current traces for the activation of TASK-3/TASK-1 by 10 μM CHET3 and blockade of 0.5 μM PK-THPP. (**B** to **G**) Representative current traces for several other K2P channels as well as other types of ion channels in response to 10 μM CHET3. CHET3 did not show strong effects on these channels. (B) Several other K2P channels. (C) hERG channel. hERG inhibitor cisapride (10 nM) was used as a reference. (D) Kv2.1 channel. Kv2.1 inhibitor TEA (50 mM) was used as a reference. (E) BK channel. BK inhibitor TEA (10 mM) was used as a reference. (F) TRPM8 channel. TRPM8 activator menthol (500 μM) was used as a reference. (G) TRPV1 channel. TRPV1 activator capsaicin (0.1 μM) was used as a reference.

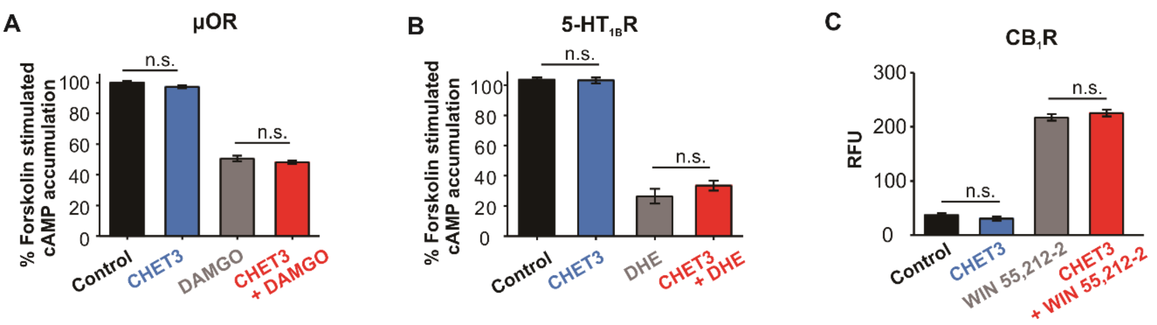


**Fig. S4. 10 μM CHET3 does not show effect on three pain-related GPCRs.** (**A**) CHET3 showed no significant effects on forskolin-stimulated cAMP accumulation in HEK293-μOR cells. CHET3 also failed to antagonize DAMGO activation on μOR (n = 5; unpaired *t* test). DAMGO is a prototypical unbiased opioid agonist. (**B**) CHET3 showed no significant effects on forskolin-stimulated cAMP accumulation in CHO-K1-5-$HT_{1B}R$ stable cells. CHET3 also failed to antagonize dihydroergotamine (DHE) activation on 5-$HT_{1B}R$ (n = 6; unpaired *t* test). DHE is a potent 5-$HT_{1B}R$ agonist. (**C**) CHET3 showed undetectable effects or competitive inhibition of WIN 55,212-2 in the calcium flux assay of CHO-$CB_1R$-Gα16 stable cells (n = 5; unpaired *t* test). WIN 55,212-2 is a potent $CB_1R$ agonist. RFU means Relative Fluorescence Units. The data were expressed as the means ± SEM performed in triplicate or quadruplicate. n.s., not significant.

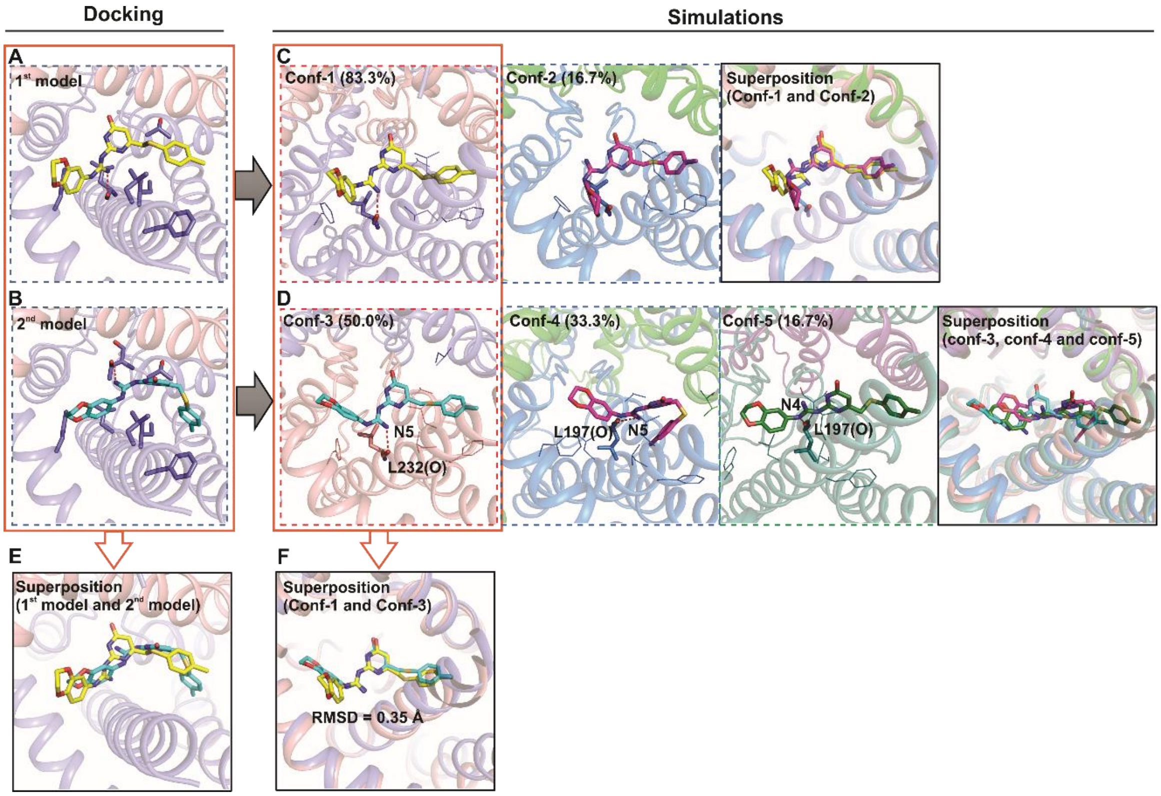


**Fig. S5. Binding modes of CHET3 suggested by docking and MD simulations.** (**A** and **B**) Docking yields two binding modes, a predominant binding mode ($1^{st}$ model, (A)) and an alternative binding mode ($2^{nd}$ model, (B)). (**C**) Two distinct poses observed in the three MD simulations initiated from $1^{st}$ model in (A). The percentages of the particular poses were labeled in parentheses. (**D**) Three representative poses observed in the three MD simulations initiated from binding $2^{nd}$ model in (B). (**E**) Superposition of the two docking models. (**F**) Superposition of the two predominant poses from different MD simulations.

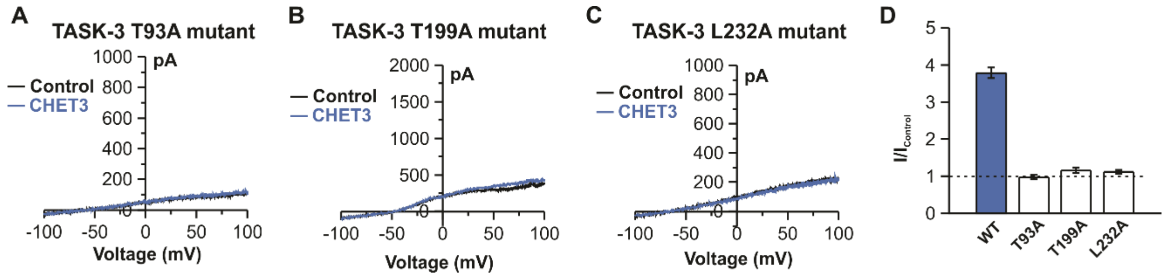


**Fig. S6. Characterization of three TASK-3 mutants.** Mutations T93A (**A**), T199A (**B**) and L232A (**C**) were no-functional channels with or without CHET3. (D) Summary for CHET3 on the three mutants (n = 9 for each mutant).

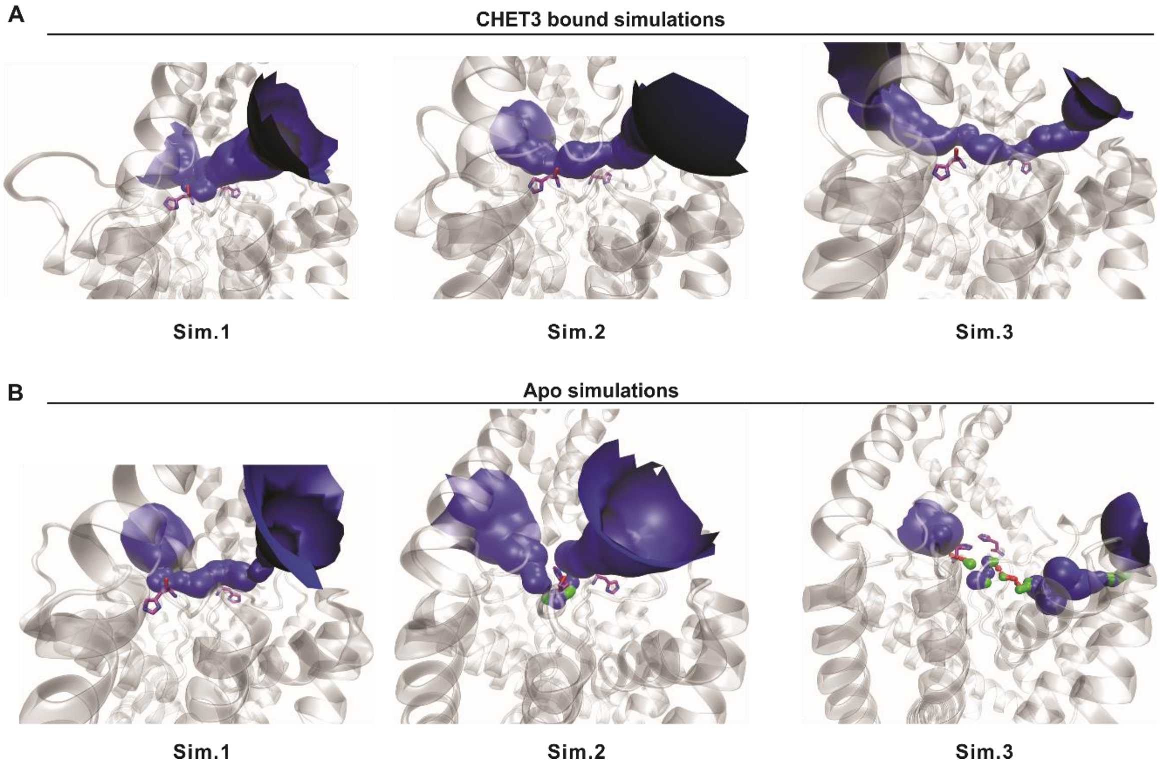


**Fig. S7. Conformations of the extracellular ion pathway in MD simulations.** (**A**) The CHET3-bound TASK-3. (**B**) The *apo* TASK-3. HOLE color code is used: blue, radius > 1.15 Å; green, radius 0.6-1.15 Å; red, radius < 0.6 Å. Residues H98 were shown in purple sticks.

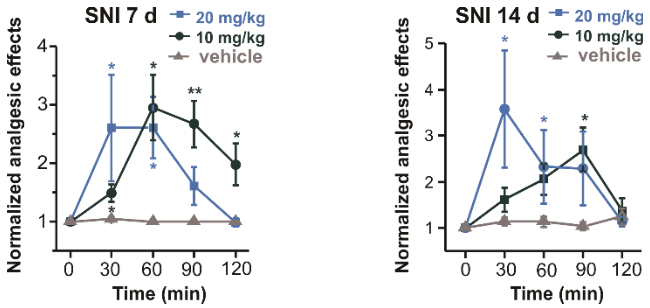


**Fig. S8. Dose-dependent analgesia by CHET3 in mechanical allodynia**. Time profile for dose-dependent analgesia by CHET3 in mechanical allodynia in rats (n = 9-13; paired sample Wilcoxon signed rank test and paired *t* test). Data are shown as mean ± SEM. *$P$ < 0.05, **$P$ < 0.01.

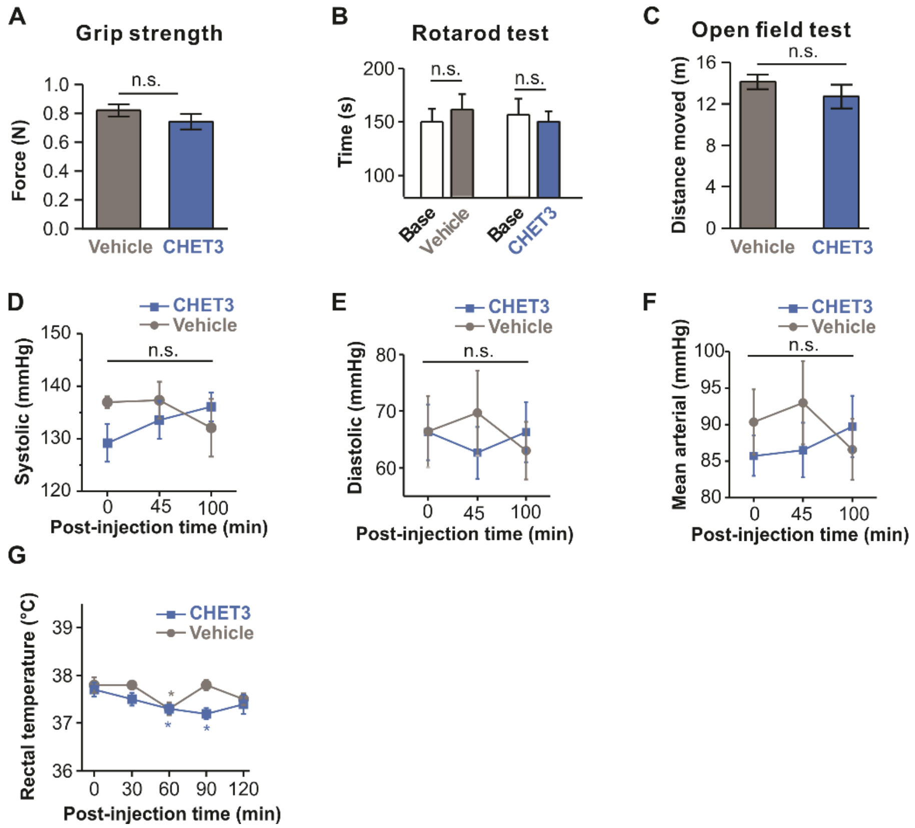


**Fig. S9. Effects of CHET3 on the locomotion activities, blood pressure and body temperature in rodents.** (**A** to **C**) Performance on locomotion activities. (A) Grip strength test in mice treated with vehicle or with CHET3 (n = 8 for vehicle, n = 9 for CHET3; unpaired *t* test). (B) Rotarod test in mice before (base) and after injection of vehicle or CHET3 (n = 10; unpaired *t* test). (C) Open field test in SNI mice (n = 9; unpaired *t* test)). (**D** to **F**) Blood pressure test in rats treated with vehicle or with CHET3 (n = 7 for vehicle, n = 8 for CHET3; D and F, paired sample Wilcoxon signed rank test; E, unpaired t test). (**G**) Rectal temperature of mice (n = 7 for vehicle, n = 8 for CHET3; paired *t* test). Data are shown as mean ± SEM. **P* < 0.05. n.s., not significant.

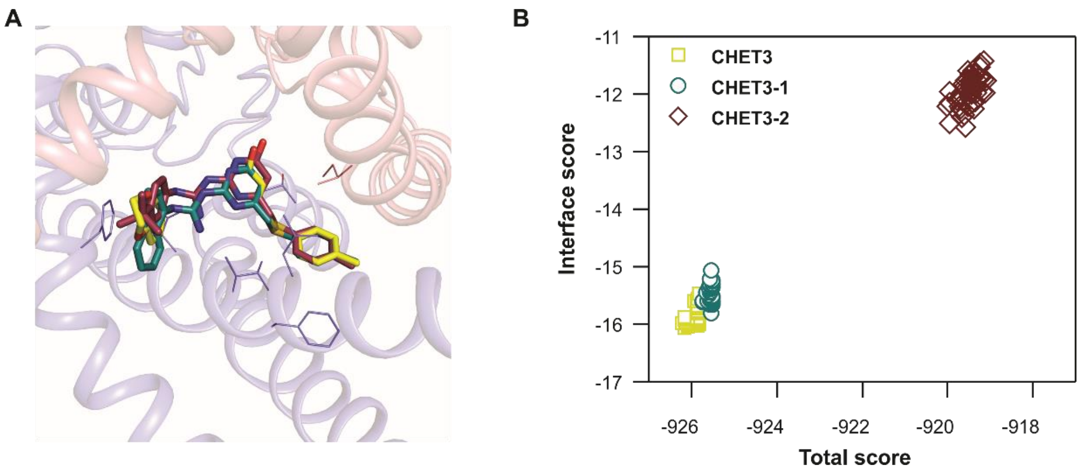


**Fig. S10. Comparison of the binding of CHET3, CHET3-1 and CHET3-2.** (**A**) Docking models comparison of CHET3 (yellow), CHET3-1 (dark cyan) and CHET3-2 (wine). Key residues around molecules were shown in lines. (**B**) Docking total scores and interface scores of CHET3 (yellow), CHET3-1 (dark cyan) and CHET3-2 (wine).

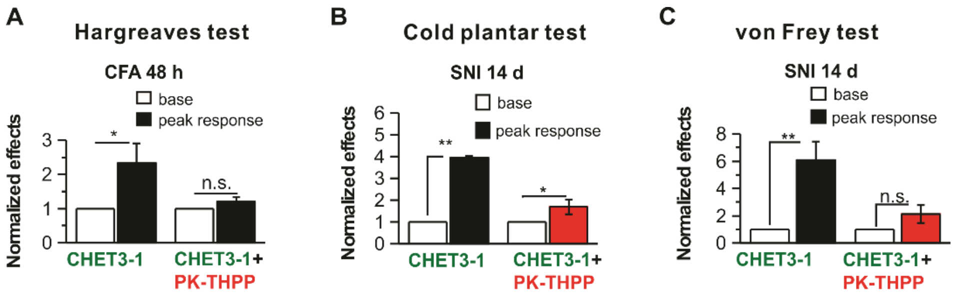


**Fig. S11. Blockade of CHET3-1 analgesia by PK-THPP.** Summary of PK-THPP on CHET3-1 analgesia in Hargreaves test (**A**), cold plantar test (**B**) and von Frey test (**C**) (n = 7-10; paired *t* test). Data are shown as mean ± SEM. $*P < 0.05$, $**P < 0.01$. n.s., not significant.

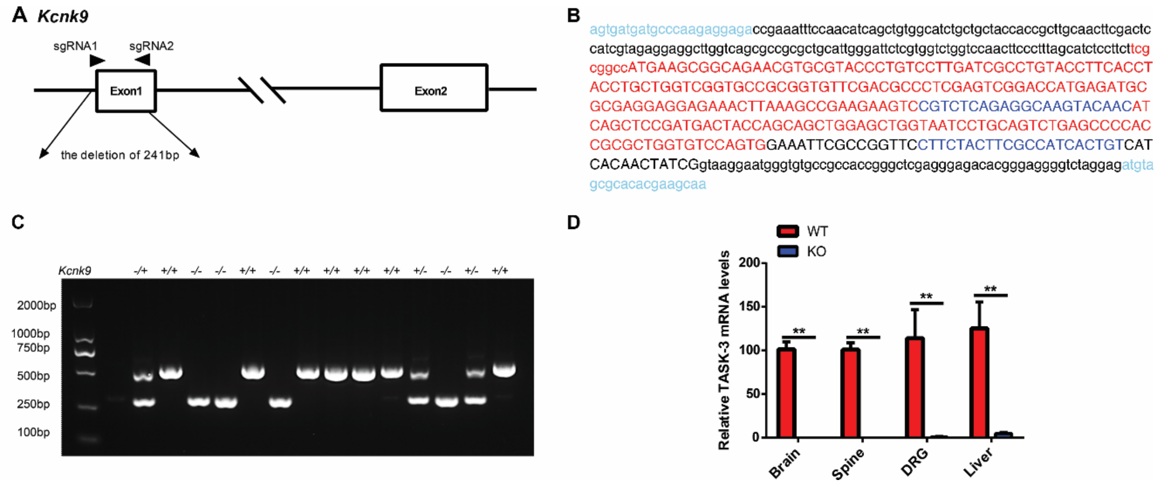


**Fig. S12. Generation and characterization of TASK-3 gene (*Kcnk9*) knockout mice.** (**A**) 241bp deletion in TASK-3 gene by CRISPR-Cas9 system. (**B**) The whole sequence of 531bp of PCR products for *Kcnk9* in WT mice with important sites highlighted by font color for characterization. The exon1 is displayed in uppercase and parts of introns flanking it are shown in lowercase. (**C**) Genotypes of WT, *Kcnk9*$^{+/-}$ and *Kcnk9*$^{-/-}$ mice were determined by PCR (The primer sequences: Forward: AGTGATGATGCCCAAGAGGAGA, Reverse: TTGCTTCGTCTCGGCTAGAT). (**D**) The relative TASK-3 mRNA expression in representative tissues in *Kcnk9*$^{-/-}$ mice compared to WT mice by qPCR (The primer sequences: Forward: CGTCTCAGAGGCAAGTACAAC, Reverse: ACAGTGTGGCGAAGTAGAAG) (n = 4; two-way ANOVA). Data are shown as mean ± SEM. **$P < 0.01$.

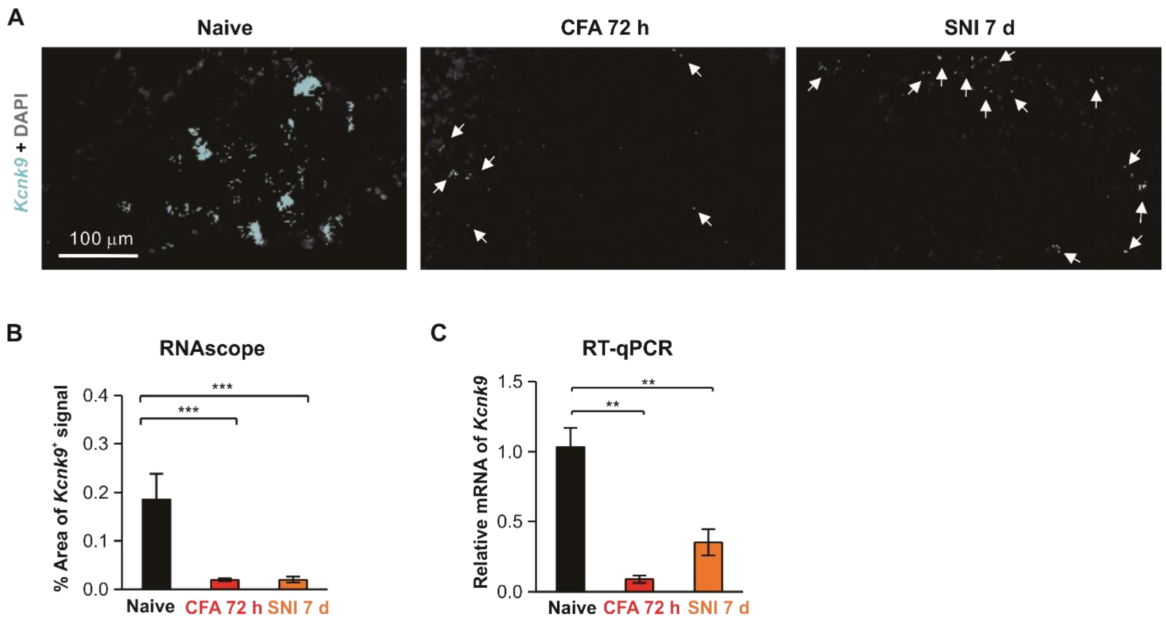


**Fig. S13. Down-regulation of peripheral TASK-3 under chronic pain.** (**A**) Images showing the down-regulation of *Kcnk9* expression in chronic pain models. (**B** and **C**) Quantification of the down-regulation of *Kcnk9* expression in chronic pain models using RNAscope (n = 11-35 sections, 3-8 mice per group; unpaired *t* test) and RT-qPCR (n = 3-5 mice per group; unpaired *t* test). Data are shown as mean ± SEM. $^{**}P < 0.01$, $^{***}P < 0.001$.

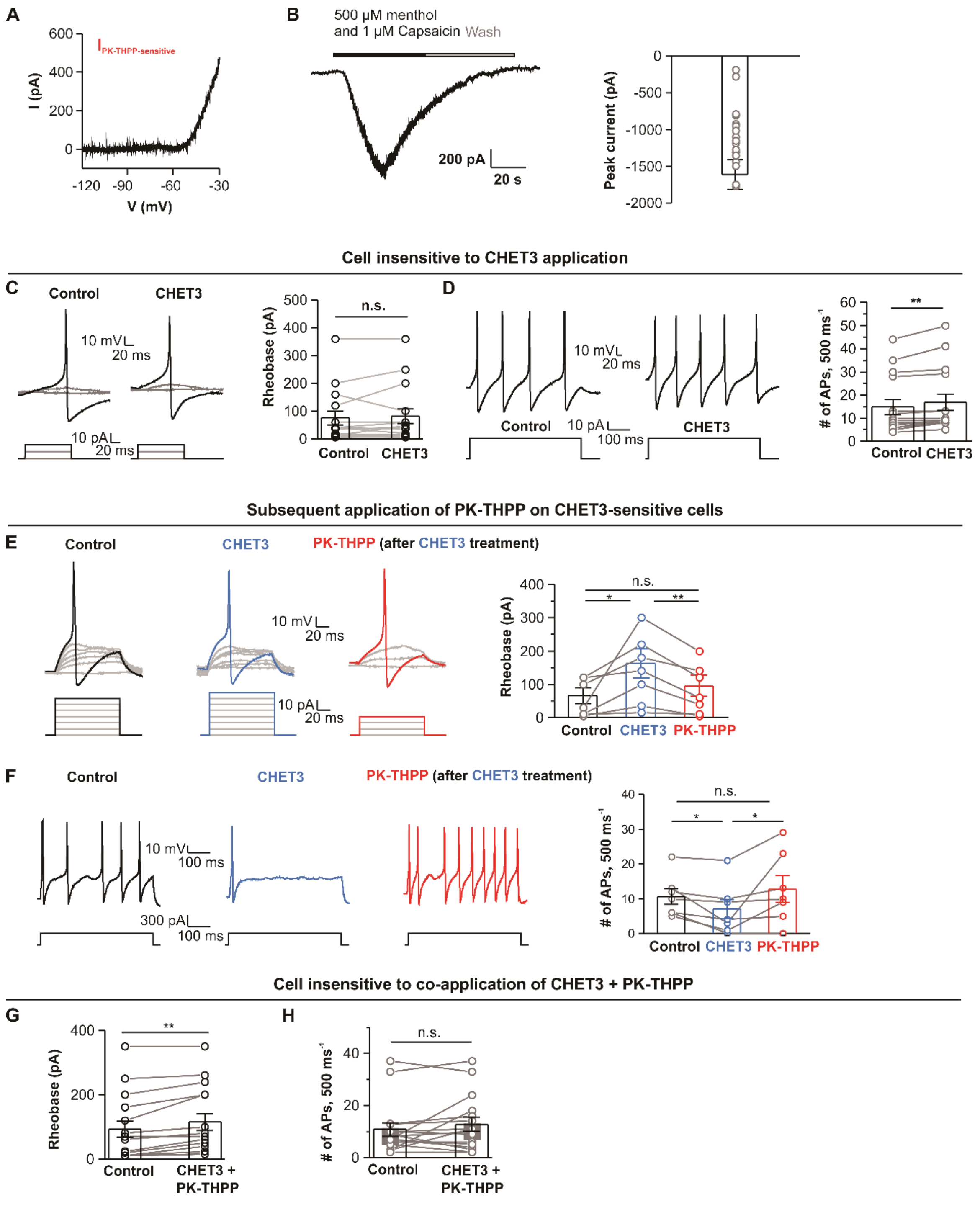


**Fig. S14. Effects of CHET3 and PK-THPP on nociceptive neurons.** (**A**) A representative trace showing PK-THPP-sensitive currents. (**B**) Trace and bar graph showing currents induced by menthol and capsaicin activation of nociceptive neurons ($V_{hold}$ of -60 mV, n =

11 cells in 5 mice). (**C** and **D**) Traces and bar graphs showing CHET3 effects on rheobase (C) and firing frequency (D) in CHET3-insensitive cells (n = 15 cells in 5 mice; paired sample Wilcoxon signed rank test in both (C) and (D)). (**E** and **F**) Traces and bar graphs showing subsequent application of PK-THPP reversed effects induced by CHET3 (n = 7 cells in 5 mice; paired sample Wilcoxon signed rank test between Control and CHET3 in (E) and (F), paired *t* test between CHET3 and PK-THPP in (E) and (F)). (**G** and **H**). Bar graphs showing the rheobase and frequency changes in cells insensitive to the co-application of CHET3 and PK-THPP (n = 16 cells in 3 mice; paired sample Wilcoxon signed rank test in both (G) and (H)).Data are shown as mean ± SEM. *$P < 0.05$, **$P < 0.01$. n.s., not significant.

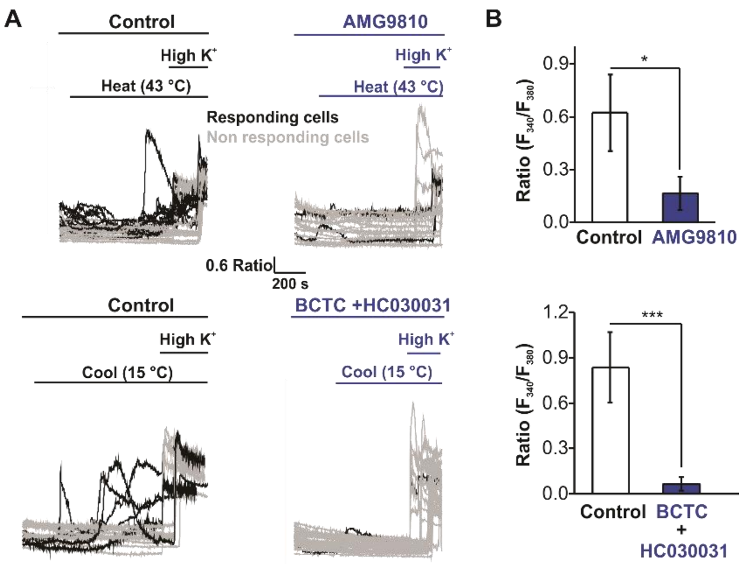


**Fig. S15. Thermal stimulation induced $Ca^{2+}$ signals were mediated by TRP channels.** (**A**) Individual $Ca^{2+}$ imaging traces from small-sized DRG neurons in representative field of views in response to heat (25 ℃-43 ℃), cooling (37 ℃-15 ℃). 5 µM AMG9810 (TRPV1 antagonist), 10 µM BCTC (TRPM8 antagonist) and 20 µM HC030031 (TRPA1 blocker) were used. (**B**) Bar graphs summary for experiments in (A). (Heat (AMG9810): n = 48 cells in 7 coverslips for control, n = 67 cells in 11 coverslips for AMG9810; Mann-Whitney test. Cool (BCTC + HC030031): n = 49 cells in 6 coverslips for control, n = 77 cells in 11 coverslips for BCTC + HC030031; Mann-Whitney test. Experiments were from 2-3 independent preparations from 4-6 mice). Data are shown as mean ± SEM. *$P < 0.05$, ***$P < 0.001$. n.s., not significant.

**Table S1. Echocardiographic evaluation of CHET3 on mice.**

| Measurement | CHET3 | Vehicle | P value |
|---|---|---|---|
| Heart Rate (bpm) | 465.2 ± 21.0 | 460.9 ± 25.7 | 0.7 |
| End-systolic left ventricular diameter (sec, mm) | 2.3 ± 0.2 | 2.2 ± 0.5 | 0.7 |
| End-diastolic left ventricular diameter (day, mm) | 3.3 ± 0.2 | 3.2 ± 0.6 | 0.6 |
| Left ventricular end-systolic volume (sec, µl) | 17.8 ± 3.9 | 17.3 ± 8.9 | 0.9 |
| Left ventricular end-diastolic volume (day, µl) | 45.2 ± 7.4 | 43.1 ± 18.2 | 0.8 |
| Stroke Volume (µl) | 27.4 ± 5.3 | 25.8 ± 9.7 | 0.7 |
| Ejection Fraction (EF, %) | 60.6 ± 6.6 | 61.5 ± 5.8 | 0.8 |
| Fractional Shortening (FS, %) | 31.6 ± 4.7 | 32.1 ± 3.7 | 0.9 |
| Cardiac Output (mL/min) | 12.7 ± 2.2 | 12.1 ± 4.9 | 0.8 |
| End-systolic left ventricular anterior wall thickness (sec, mm) | 1.6 ± 0.2 | 1.5 ± 0.2 | 0.7 |
| End-diastolic left ventricular anterior wall thickness (day, mm) | 1.1 ± 0.3 | 1.1 ± 0.2 | 0.9 |
| End-systolic left ventricular posterior wall thickness (sec, mm) | 1.2 ± 0.2 | 1.4 ± 0.4 | 0.4 |
| End-diastolic left ventricular posterior wall thickness (day, mm) | 0.9 ± 0.1 | 1.1 ± 0.4 | 0.4 |

Data are shown as mean ± SD. n = 7 for vehicle, n = 8 for CHET3; unpaired *t* test.

**Table S2. CHET3 pharmacokinetic in plasma and brain following a single intraperitoneal administration to naïve male C57BL/6 mice.**

| **Dose** | **i.p. 10 mg/kg** | |
|---|---|---|
| **Tissue** | **Plasma** | **Brain** |
| $T_{max}$ (hour) | 0.083 | 0.25 |
| $C_{max}$ (ng/mL) | 1112.00 | 79.10 |
| $AUC_{0\text{-}last}$ (hr*ng/mL) | 489.88 | 40.77 |
| $T_{1/2}$ (hour) | 3.11 | 2.42 |
| Tissue/plasma AUC ratio | NA | 0.08 |

Abbreviations: $T_{max}$, time to $C_{max}$; $C_{max}$, maximum plasma concentration; AUC, area under the plasma or tissue concentration–time curve; $T_{1/2}$, terminal half-life; NA, not applicable.

**Table S3. CHET3 pharmacokinetic in plasma and brain following a single intraperitoneal administration to SNI 7-d male C57BL/6 mice.**

| Dose | i.p. 10 mg/kg | |
|---|---|---|
| **Tissue** | **Plasma** | **Brain** |
| $T_{max}$ (hour) | 0.25 | 0.25 |
| $C_{max}$ (ng/mL) | 1337.50 | 64.88 |
| $AUC_{0\text{-}last}$ (hr*ng/mL) | 979.28 | 69.91 |
| $T_{1/2}$ (hour) | 1.33 | 1.20 |
| Tissue/plasma AUC ratio | NA | 0.05 |

Abbreviations: $T_{max}$, time to $C_{max}$; $C_{max}$, maximum plasma concentration; AUC, area under the plasma or tissue concentration–time curve; $T_{1/2}$, terminal half-life; NA, not applicable.
